# The Evolution of Chunks in Sequence Learning

**DOI:** 10.1101/2021.02.12.430894

**Authors:** Laure Tosatto, Joël Fagot, Dezso Nemeth, Arnaud Rey

## Abstract

Chunking mechanisms are central to several cognitive processes and notably to the acquisition of visuo-motor sequences. Individuals segment sequences into chunks of items to perform visuo-motor tasks more fluidly, rapidly, and accurately. However, the exact dynamics of chunking processes in the case of extended practice remain unclear. Using an operant conditioning device, eighteen Guinea baboons (*Papio papio*) produced a fixed sequence of nine movements during 1,000 trials by pointing to a moving target on a touch screen. Response times analyses revealed a specific chunking pattern of the sequence for each baboon. More importantly, we found that these patterns evolved during the course of the experiment, with chunks becoming progressively fewer and longer. We identified two chunk reorganization mechanisms: the *recombination* of preexisting chunks and the *concatenation* of two distinct chunks into a single one. These results provide new evidence on chunking mechanisms in sequence learning and challenge current models of associative and statistical learning.

---

A key mechanism allowing our cognitive system to compress information and increase short term memory capacity is the formation of chunks (Mathy & Feldman, 2012; Miller, 1956). Chunking is defined as the process of associating and grouping several items together into a single processing unit (Gobet & al, 2001; Gobet, Lloyd-Kelly and Lane, 2016). In coherence with the limits encountered by our cognitive system (Cowan, 1988, 2017), the storage capacity of the chunks themselves seems limited (3 or 4 items per chunk, Allen & Coyne, 1988; Chase & Simon, 1973; Johnson, 1970). In the field of perceptual-motor learning, chunking has been considered as the main motor sequence integration mechanism (Diedrichsen & Kornysheva, 2015; Wymbs et al. 2012).

Perceptual-motor sequence learning is commonly described as the process by which a sequence of movements is acquired and executed with increased speed and accuracy (Willingham, 1998). Current sequence learning paradigms use a sequential button-press task: subjects are presented with a series of visual stimuli organized in a sequence and are asked to press a corresponding response button (e.g., Cohen, Ivry, & Keele, 1990; Nemeth et al., 2010; Verwey, 2001). A faster and more accurate performance over time reflects learning of the sequence. Typically, researchers have reported robust effects of sequence’s length on subjects’ response times (RTs): for short sequences (<4 keypresses), a decrease in the successive RTs can be observed when the sequence is learned; for longer sequences though, an initial decrease in RTs for the first 3 to 4 keypresses is followed by a longer RT on the next position. Then, RTs start decreasing again for the next 3 to 4 keypresses (e.g., Bo & Seidler, 2009; Verwey & Eikelboom, 2003). Similar results have been observed in non-human subjects (Terrace, 2002; Scarf et al., 2018). This phenomenon has been interpreted as the long sequence being spontaneously segmented into shorter motor chunks reflecting the sequence organization in memory (Sakai et al., 2003). Long temporal gaps between responses are assumed to mark chunk boundaries (Abrahamse et al., 2013; Bottary et al., 2016). Yet, few studies have been interested in the evolution of chunks in the case of extended practice (e.g., Ramkumar et al., 2016; Song & Cohen, 2014; Wymbs et al., 2012). The present study aims at collecting new evidence on the evolution of chunking mechanisms in a perceptual-motor sequence learning task.

One of the main issues in most sequence learning studies is the inconsistent transition probabilities (TPs) between items of the sequence. Indeed, in an experimental set up where only a few response buttons are available to the subject (e.g., Grafton, Hazeltine and Ivry, 2002; Verwey, 2001; Willingham, 1999), some stimuli are necessarily presented multiple times within the same sequence. For instance, in a sequence such as A-B-A-C-D-B-C, A is either followed by B or C, therefore the probability of B given A is 0.5, whereas D is always followed by B thus the probability of B given D is 1. This heterogeneity of TPs may constrain the strength of the connections formed between the successive elements of the sequence and affect the resulting chunking pattern.

Another characteristic of previous studies is, apart from a few exceptions (e.g., Conway & Christiansen, 2001), that most sequence learning studies have been conducted with human participants. However, even if implicit instructions are provided, humans almost systematically use their inner language and develop strategies to perform the task (Rey et al., 2019). This may affect both their performance and their chunking processes, and it is hard to tease apart these verbal and explicit influences from associative and chunking mechanisms.

To avoid these difficulties, we first chose to test non-human primates for neutralizing the possible effect of language-based strategies. Second, we used sequences composed of items only occurring once per sequence, i.e., with a transitional probability between elements equal to 1, using the serial pointing task proposed by Minier et al. (2016). In this task, participating monkeys had to track and touch target locations in a 3 x 3 matrix of crosses displayed on the screen. They were exposed to the same sequence of 9 locations that always appeared in the same order upon a thousand trials. RTs for each location were recorded, providing us with detailed information on the temporal dynamics of sequence processing. This extended practice on a long single repeated sequence allowed us to collect new evidence on the formation and the evolution of chunks over time.

## Method

### Participants

Thirteen female and five male Guinea baboons (*Papio papio*, age range 2.8—23.7 years) from the CNRS primate facility in Rousset (France), living in a social group of 25 individuals, were tested in this study. Water was provided *ad libitum* during the test, and the monkeys received their normal food ration of fruits every day at 5 PM. For practical reasons, we stopped the experiment after 18 monkeys completed all scheduled trials.

### Materials

#### Apparatus

This experiment was conducted with a computer-learning device based on the voluntary participation of baboons (for details, see Fagot & Bonté, 2010). Baboons implanted with a RFID microchip had free access to 10 automatic operant conditioning learning devices equipped with touch screens. Each time a monkey entered a test chamber, it was identified by its microchip, and the system resumed the trial list where the subject left it at its previous visit. The experiment was controlled by E-prime (Version 2.0, Psychology Software Tools, Pittsburgh, PA, USA).

#### Task and stimuli

The screen was divided into nine equidistant predetermined locations represented by white crosses on a black background, virtually labeled as position 1 to 9 (see Figure 1). A trial began with the presentation of a yellow fixation cross at the bottom of the screen. Once pressed, the fixation cross disappeared and the nine white crosses were displayed, one of them being replaced by the target, a red circle. When the target was touched, it was immediately replaced by the cross. The red circle then replaced the next position in the sequence until it was touched, and a new position was displayed. Reward was provided at the end of a sequence of nine touches.

**Figure 1.**
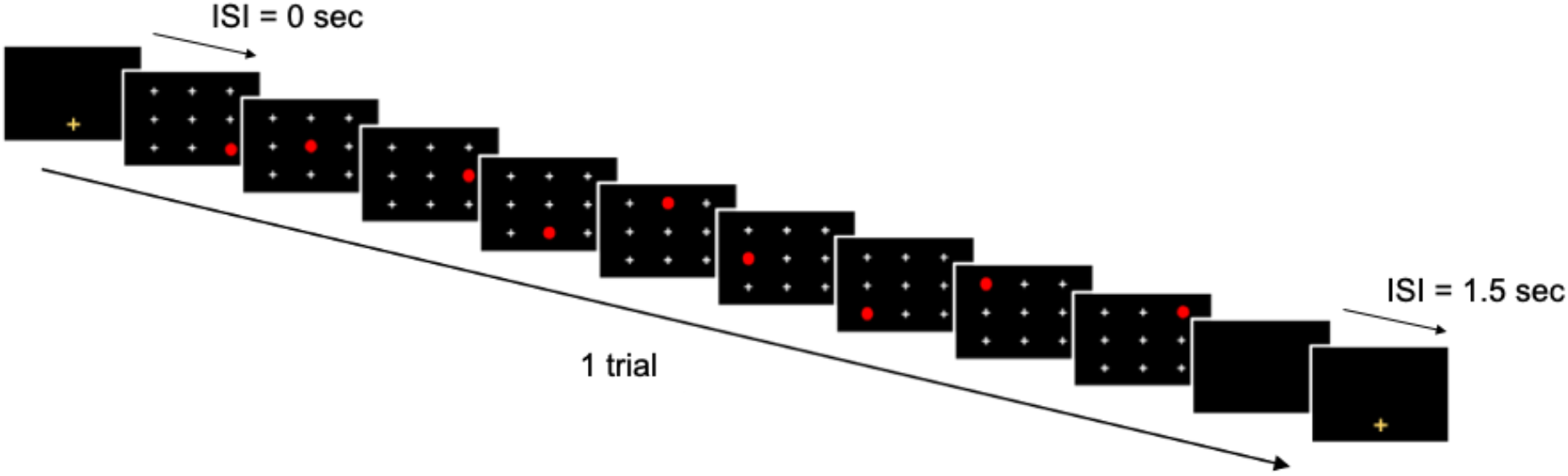
Experimental display and stimuli presentation.

To learn the task, baboons initially received random trials that were rewarded after three touches. Then, the number of touches in a trial was progressively increased up to nine. If baboons touched an inappropriate location (incorrect trial) or failed to touch the screen within 5,000ms after the red circle’s appearance (aborted trial), a green screen was displayed for 3,000ms as a marker of failure. Aborted trials were not retained and therefore presented again, while incorrect trials were not. The time elapsed between the appearance of the red circle and the baboon’s touch on this circle was recorded as the RT.

#### Design of the sequences

To control the motor difficulty of the transitions to be produced in the sequence, a random phase of sequence production was first conducted, where thirteen baboons performed random sequences of six positions for 1,000 trials. Based on these random trials, a baseline measure for all possible transitions from one location to another was computed by calculating mean RTs for each transition (e.g., from position 1 to 9), leading to a 9×9 matrix of mean RTs calculated over the entire group of baboons (see Appendix A).

Based on these baseline measures, we designed two sequences of nine serial positions for which each transition T was faster (or equally fast) to produce than the next one (i.e., T1≤T2≤…≤T8; see Appendix B). This way, a decrease in RT for a given transition can be interpreted as the anticipation (or learning) of that position from the previous one.

### Procedure

Baboons were either presented with Sequence 1 (N=8) or Sequence 2 (N=10) and had to produce it for 1000 successive trials. RTs for each position of the sequence were recorded for all the trials.

## Results

On average, baboons required 2.9 days to complete the 1,000 trials, with a mean of 339.7 trials per day and a mean accuracy level of 92.5% (*SD*=6.2%). Incorrect trials were removed from the dataset (7.8%). A recursive trimming procedure excluded RTs greater or smaller than 2.5 standard deviations from the subject’s mean for each of the nine possible positions (24.4%)^1^. RTs for each of the nine positions and for the 1,000 trials were divided into 10 Blocks of 100 trials.

### General sequence learning

Learning of the sequence was estimated on mean RTs by a repeated measures one-way ANOVA with Block (1-10) as the within factor. The effect of Block was highly significant (Block 1, *M*= 452.8, *SD*=45.3; Block 10, *M*=400.1, *SD*=56.3), *F*(1,9)=30.43, *p*<.001, *η^2^*=0.642, indicating that mean RTs decreased throughout the blocks of trials and that monkeys learned the sequence.

### Chunking of the sequence

To study the chunking pattern of the sequence by monkeys, we adopted the following reasoning. We considered successive positions *A* and *B* to be part of the same chunk as long as the transition time from one position to the next did not correspond to a significant increase in RT. Therefore, if RT_A_<RT_B_ then *A* and *B* do not belong to the same chunk, and the AB transition marks a chunk boundary (Kennerly et al., 2004). This reasoning was applied on each Block and for each monkey. Statistical significance was assessed through paired-sample t-tests for each pair of successive positions. Each time the RT of a pair’s second position was significantly higher than the first position, it marked a chunk boundary. Figure 2 illustrates this method on the performances of one monkey.

**Figure 2.**
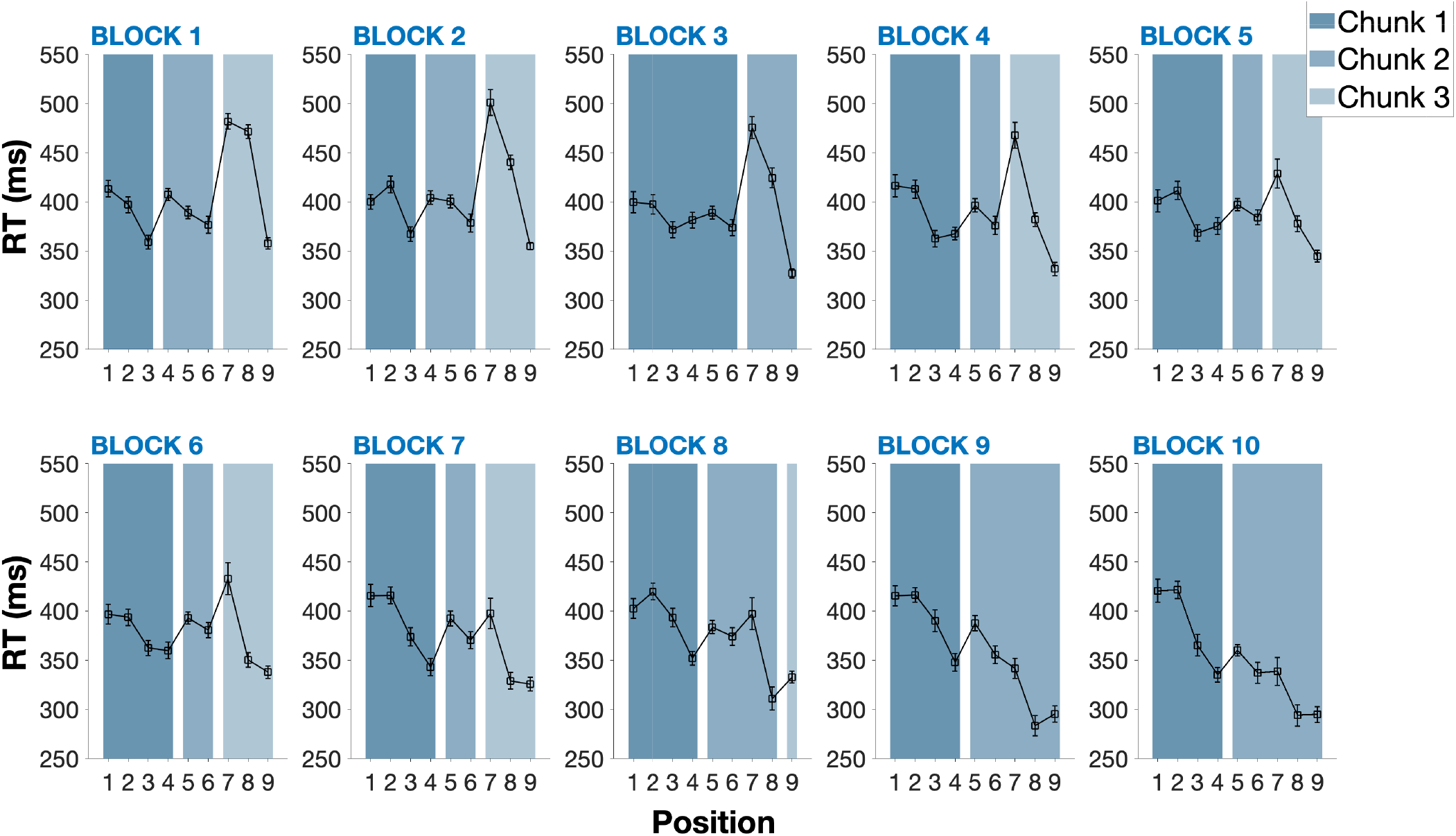
Evolution of the chunking pattern for one individual (Atmosphere) throughout the task. *Note*. Mean RT per position across the 10 blocks of trials for one baboon (Atmosphere) showing the evolution of the chunking pattern. This individual initially parses the sequence into three chunks of three positions in the first three blocks of trials. A reorganization starts in Block 4, as Position 4 is integrated into the first chunk. Another reorganization occurs in Block 8, where the 2^nd^ and 3^rd^ chunks are progressively concatenated (error bars represent 95% confidence intervals).

### Evolution of chunks

With this method, we were able to quantify the number of chunks and their average size produced on each block by each monkey. For example, the monkey from Figure 2 produced 3 chunks and an average chunk size of 3 in Block 1. In Block 10, the number of chunks was 2 and the average chunk size was 4.5. Figure 3 reports the average number of chunks and chunk size for all monkeys across the 10 blocks of the experiment. For example, we found that the mean chunk size was 2.2 positions (*CI*=0.31; *Min*=1; *Max*=5) in the first block and 3.375 (*CI*=0.53; *Min*=1; *Max*=8) in the last block. Two repeated measures one-way ANOVA were conducted to test the effect of Block on the mean number of chunks and the average chunk size. This analysis revealed that the number of chunks significantly decreased across blocks, *F*(1,9)=9.421, *p*<.001, *η^2^*=.357, and that the average chunk size significantly increased across blocks, *F*(1,9)=4.794, *p*<.01, *η^2^*=22.

**Figure 3.**
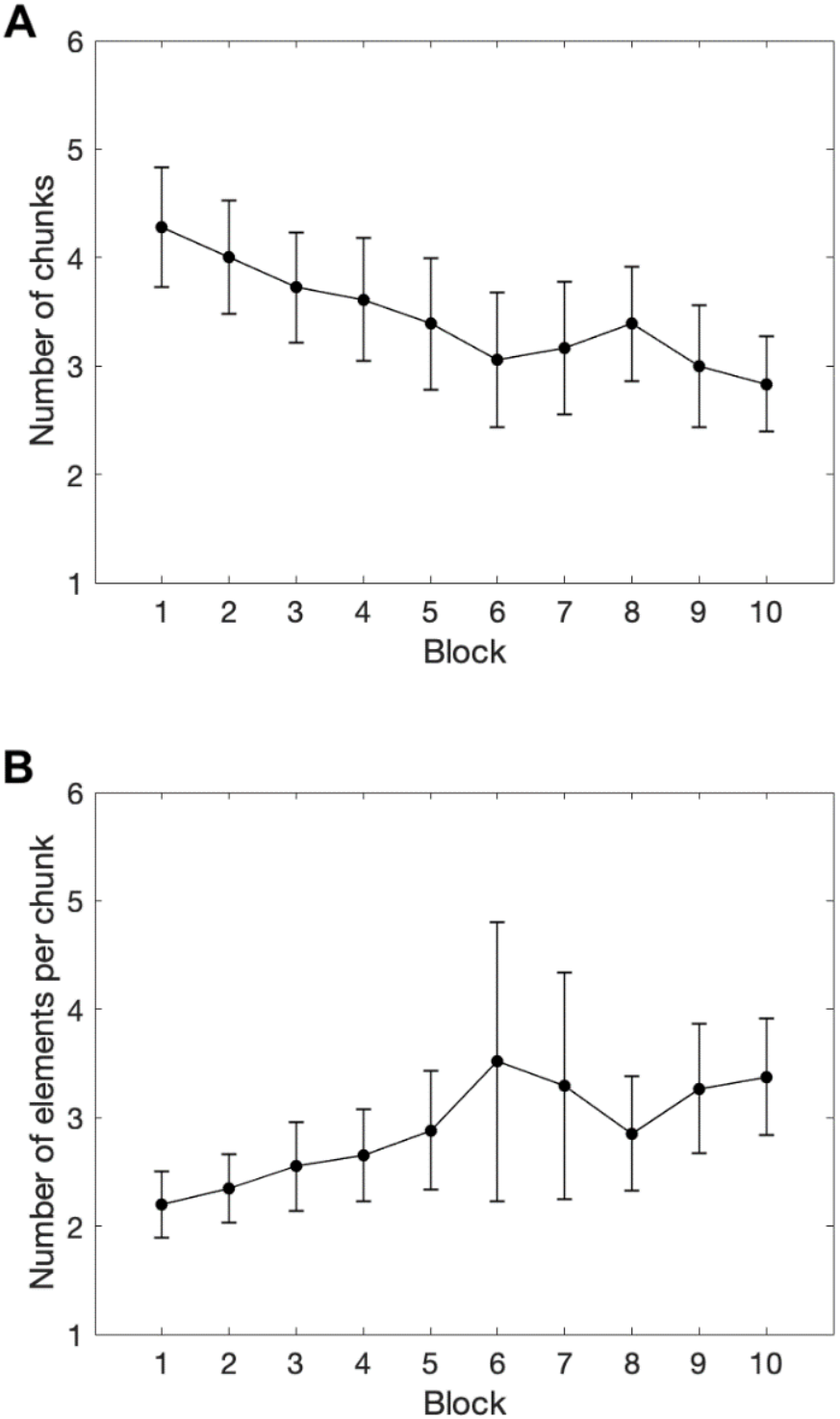
Evolution of chunks across blocks. *Note*. A. Mean number of chunks per block. B. Mean chunk size (i.e. number of elements per chunk) per block (error bars represent 95%confidence intervals).

The evolution of these two indicators can be accounted for by two reorganization mechanisms: the *recombination* of pre-existing chunks (50.4% of the reorganizations, observed for 15 monkeys) and the *concatenation* of two chunks into a single one (49.6% of the reorganizations, observed for 17 monkeys). An illustration of these reorganizations is provided in Figure 2: from Block 1 to Block 3, the first six positions of the sequence were parsed into two equivalent chunks of three positions, indicated by the significant increase in RTs between Position 3 and 4. Starting Block 4, the chunks are recombined with Position 4 being slowly integrated to the first chunk as its difference with Position 3 disappears and a significant increase in RTs is now appearing between Position 4 and 5. The concatenation of two chunks into a single one is also illustrated in Figure 2: while Chunks 2 and 3 were separated by a significant increase in RT between Position 6 and Position 7 until Block 8, this difference disappeared in Block 9, and the two chunks were then grouped into one. Table 1 provides a detailed description of the chunks appearing in Figure 2. For each block of trials, it counts the number of chunks, their respective size, and each occurrence of a concatenation or a recombination. At the group level, Table 2 provides the total number of concatenations and recombinations obtained for each block and for all monkeys^2^. A repeated measure ANOVA with Block and Mechanisms (concatenation vs. recombination) did not reveal any significant effect (all ps > .05).

**Table 1.** Summary table of the reorganizations observed throughout the task for one baboon (Atmophere).

|  | Chunk size |  |  | Nb of chunks | Concatenation | Recombinaison |
| --- | --- | --- | --- | --- | --- | --- |
|  | C1 | C2 | C3 |  |  |  |
| Block 1 | 3 | 3 | 3 | 3 | - | - |
| 2 | 3 | 3 | 3 | 3 | - | - |
| 3 | 3 | 3 | 3 | 3 | - | - |
| 4 | 4 | 2 | 3 | 3 | - | 1 |
| 5 | 4 | 2 | 3 | 3 | - | 1 |
| 6 | 4 | 2 | 3 | 3 | - | - |
| 7 | 4 | 2 | 3 | 3 | - | - |
| 8 | 4 | 4 | 1 | 3 | - | 1 |
| 9 | 4 | 5 | - | 2 | 1 | - |
| 10 | 4 | 5 | - | 2 | - | - |
| Total |  |  |  |  | 1 | 3 |

*Note.* Numbers in columns 2-4 refer respectively to the size of the first (C1), second (C2) and third (C3) chunk (if any) in the sequence.

**Table 2.** Total number of concatenations and recombinations per block.

| Block | Concatenations | Recombinations |
| --- | --- | --- |
| 2 | 11 | 9 |
| 3 | 8 | 7 |
| 4 | 6 | 7 |
| 5 | 6 | 4 |
| 6 | 8 | 5 |
| 7 | 3 | 6 |
| 8 | 1 | 10 |
| 9 | 8 | 3 |
| 10 | 5 | 6 |
| Total | 56 | 57 |

### Additional analyses

Our data show that a chunking pattern emerges and evolves over time for each monkey (i.e., Figures S1 to S17). The averaged data on Block 1 indicates that a first chunking pattern rapidly emerges after a few dozen of trials. However, we checked that monkeys produced the same ascending pattern of RTs during their first 10 trials on the repeated sequence than the one we computed from the random baseline trials (see Figure S18). This ensures that our sequences were correctly designed and that chunking mechanisms are rapidly efficient.

We also compared the mean response times computed on every trial and averaged over blocks in the random and repeated sequence conditions. The result indicates that chunking mechanisms qualitatively and quantitatively influenced the monkey’s behavior by significantly increasing their global efficacy in this visuo-motor task (see Figure S19).

## Discussion

Three main findings were obtained in the present study. First, we found that non-human primates spontaneously segmented long sequences into short chunks. Second, with extended practice, chunks became longer and fewer. Third, based on these observations, we assumed this decrease in the number of chunks and this increase in chunks’ size was due to two types of reorganizations: the recombination of several preexisting chunks and the concatenation of two distinct chunks into a single one.

Our first finding is consistent with previous studies on sequence learning in humans as participants were found to initially segment the sequence into small chunks (e.g., Nissen & Bullemer, 1987; Verwey, 1996; 2001; 2003, Verwey et al., 2002). Chunks typically contained 2 to 4 items, sometimes 5 (Sakai et al., 2003), which is similar to what we observed in the first block of the experiment. With extended practice, the mean chunk size could reach up to 8 successive positions. However, these large chunks are very rare in our data and in the literature, although Kennerly, Sakai and Rushworth (2004) found in humans a mean chunk size of 7.83 items for sequences of 12 items. This difference is likely due to motor constrains that vary across experimental paradigms and may facilitate the development of longer chunks. This first result indicates that chunking is a fundamental mechanism of sequence learning in both human and non-human primates, and that the cognitive system spontaneously forms strong associations between repeated co-occurring events (e.g., Perruchet & Vinter, 1998; 2002; Rey et al., 2019). However, the results also show that there is an initial limit to the number of associations that can be formed successively.

Our second finding indicates that extended practice allows the baboons to exceed these limits, as we found that chunks can be reorganized into longer segments. This feature is not systematic as we observed reorganizations in 17 monkeys out of 18. Ramkumar et al. (2016) suggested that to limit the cost of computation, learning new sequences of movements starts with many short chunks. With practice, the execution of short chunks becomes more efficient, which reduces the computation’s complexity. This increase in efficiency for short chunks would promote more complex computations leading to the development of longer chunks. However, the precise neural mechanisms supporting these changes over time are still unclear, as are the conditions that promote switching from one pattern to another.

Our third main finding is related to the evolution of chunks and the two reorganization mechanisms that allow chunks to become longer: the concatenation and recombination of chunks. Abrahamse et al. (2013) define the concatenation mechanism as the process by which two successive chunks are performed more fluidly and the temporal gap between them decreases. This description is consistent with our findings but, to our knowledge, there is no other report of two successive chunks becoming one and the temporal gap between them disappearing. As for recombinations, they correspond to the emergence of a new segmentation pattern across chunks. As shown in Table 2, the occurrence of concatenation or recombination does not seem related to a specific stage of learning. In both cases, one can assume that these modifications in the chunking pattern is certainly related to the increase in efficiency in the realization of some chunks (Ramkumar et al., 2026). This would then modify the stability of the preexisting chunking pattern and favor either the concatenation of two chunks into a longer one or local rearrangements leading to recombinations and to more functional chunking patterns.

Interestingly, current models of chunking only partially account for the reorganizations we found. Most models, like the Competitive Chunking model (Servan-Schreiber & Anderson, 1990) or the PARSER model (Perruchet & Vinter, 1998) assume that repeated sequences lead to the formation of chunks that are stored in memory. They also assume that two co-occurring chunks can lead to a new and larger chunk by concatenating two previously smaller chunks. Recombination, though, is not a mechanism implemented in any model of chunking. The present data, therefore, represent a strong challenge for current models of statistical and associative learning.

To summarize, these data indicate that when non-verbal non-human animals are repeatedly exposed to a long sequence of 9 elements, associations are formed initially between a limited number of co-occurring elements (i.e., from 2 to 4 elements). With extended practice, these patterns of associations can become longer. The reorganization mechanisms supporting these evolutions are not yet accounted for by any model of associative learning but understanding the dynamics of these mechanisms represents a strong challenge for the future generation of computational models of associative and statistical learning.

## Context

We continuously learn and encode the statistical regularities appearing in our environment. Previous empirical and theoretical studies have proposed that chunking and associative learning mechanisms are good candidates to account for our ability to encode sequential regularities. We were interested in the present study in determining the dynamics of these chunking mechanisms when the sequential information was composed of a long, repeated sequence. This sequence was processed by a non-human primate species that does not have access to language recoding abilities. Consistent with previous theoretical accounts, we found that the long sequence was initially parsed into small chunks. After extensive training, some of these small chunks were concatenated leading to longer chunks and to a more efficient processing of the sequence. However, we also found other types of chunk reorganizations, that we called “recombinations”, that are not predicted by current models of statistical and associative learning. These results will certainly provide new constraints for elaborating the next generations of computational models accounting for chunking mechanisms.

## Acknowledgements

This work was supported by the BLRI Labex (ANR-11-LABX-0036), Institut Convergence ILCB (ANR-16-CONV-0002), the CHUNKED ANR project (#ANR-17-CE28-0013-02) and IDEXLYON Fellowship of the University of Lyon as part of the Programme Investissements d’Avenir (ANR-16-IDEX-0005). The funders had no role in study design, data collection and analysis, decision to publish, or preparation of the manuscript.We are grateful to Laura Ordonnez-Magro for her helpful comments on a previous version of the manuscript.

## Open Practices Statements

Data from the experiment are available on Open Science Framework at https://osf.io/xcw95/.

## Animal rights

This research adhered to the applicable French rules for ethical treatment of research animals and received ethical approval from the French Ministry of Education (approval APAFIS#2717-2015111708173794 10 v3).

## Appendix A

Mean response times over the entire group of baboons for each of the 72 possible transitions calculated from the 1000 random trials.

| 1<sup>st</sup> position in Transition | 2<sup>nd</sup> position in Transition |  |  |  |  |  |  |  |  |
| --- | --- | --- | --- | --- | --- | --- | --- | --- | --- |
|  | 1 | 2 | 3 | 4 | 5 | 6 | 7 | 8 | 9 |
| 1 | - | 519 | 573 | 495 | 482 | 509 | 521 | 497 | 543 |
| 2 | 569 | - | 553 | 513 | 474 | 511 | 523 | 491 | 509 |
| 3 | 558 | 519 | - | 513 | 472 | 488 | 544 | 493 | 512 |
| 4 | 551 | 517 | 560 | - | 464 | 509 | 522 | 482 | 546 |
| 5 | 549 | 504 | 552 | 501 | - | 483 | 535 | 479 | 527 |
| 6 | 567 | 515 | 546 | 507 | 484 | - | 533 | 483 | 511 |
| 7 | 555 | 504 | 558 | 475 | 463 | 516 | - | 484 | 541 |
| 8 | 554 | 512 | 540 | 485 | 448 | 472 | 512 | - | 507 |
| 9 | 546 | 512 | 540 | 514 | 460 | 464 | 550 | 485 | - |

*Note.* All transitions are in milliseconds (ms) and correspond to the time elapsed between the disappearance of the red circle from the 1<sup>st</sup> position of the Transition and the monkey's touch on the 2<sup>nd</sup> position of the Transition.

## Appendix B

Selected sequences and corresponding mean transition times (based on the baseline acquisition, see Appendix A)

Appendix B *Selected sequences and corresponding mean transition times (based on the baseline acquisition, see Appendix A)*
| Sequence | Position |  |  |  |  |  |  |  |  | Mean transition time (ms) |  |  |  |  |  |  |  |
| --- | --- | --- | --- | --- | --- | --- | --- | --- | --- | --- | --- | --- | --- | --- | --- | --- | --- |
|  | 1 | 2 | 3 | 4 | 5 | 6 | 7 | 8 | 9 | T1 | T2 | T3 | T4 | T5 | T6 | T7 | T8 |
| Seq. 1<br>(N=8) | 9 | 5 | 6 | 8 | 2 | 4 | 7 | 1 | 3 | 460 | 483 | 483 | 512 | 513 | 522 | 555 | 573 |
| Seq. 2<br>(N=10) | 7 | 5 | 6 | 8 | 2 | 4 | 9 | 1 | 3 | 463 | 483 | 483 | 512 | 513 | 522 | 555 | 573 |

## Footnotes

1 Inspection of the response times distribution revealed that a majority of responses were produced around 500ms. A smaller group of RTs appeared around 1,000 ms and was likely due to situations in which baboon’s response was not recorded by the computer, because their hands were dirty. In this situation, they had to touch the screen again, and longer RTs were recorded (that are on average twice longer compared to the first responses). This is why we have adopted this recursive trimming procedure. After applying this procedure, there was still a mean number of 77.9 remaining RTs for each position per participant and per block of 100 trials (the minimum number of remaining RTs for one position being 32).

2 See Supplementary material for a description of the patterns of responses for each monkey over blocks. The same information as the one displayed in Figure 2 and Table 1 for the monkey “Atmosphere” is provided for all monkeys from the present experiment (see Figures and Tables S1 to S17). These data clearly show that each monkey displays a specific chunking pattern that follows a specific evolution over blocks of trials.

