## Supplementary Materials for "The Evolution of Chunks in Sequence Learning"

#### Figure S1-S17 and Table S1-S17: Subject by subject chunking pattern performance

#### Figure and Table S1

*Evolution of the chunking pattern for Angèle*

A

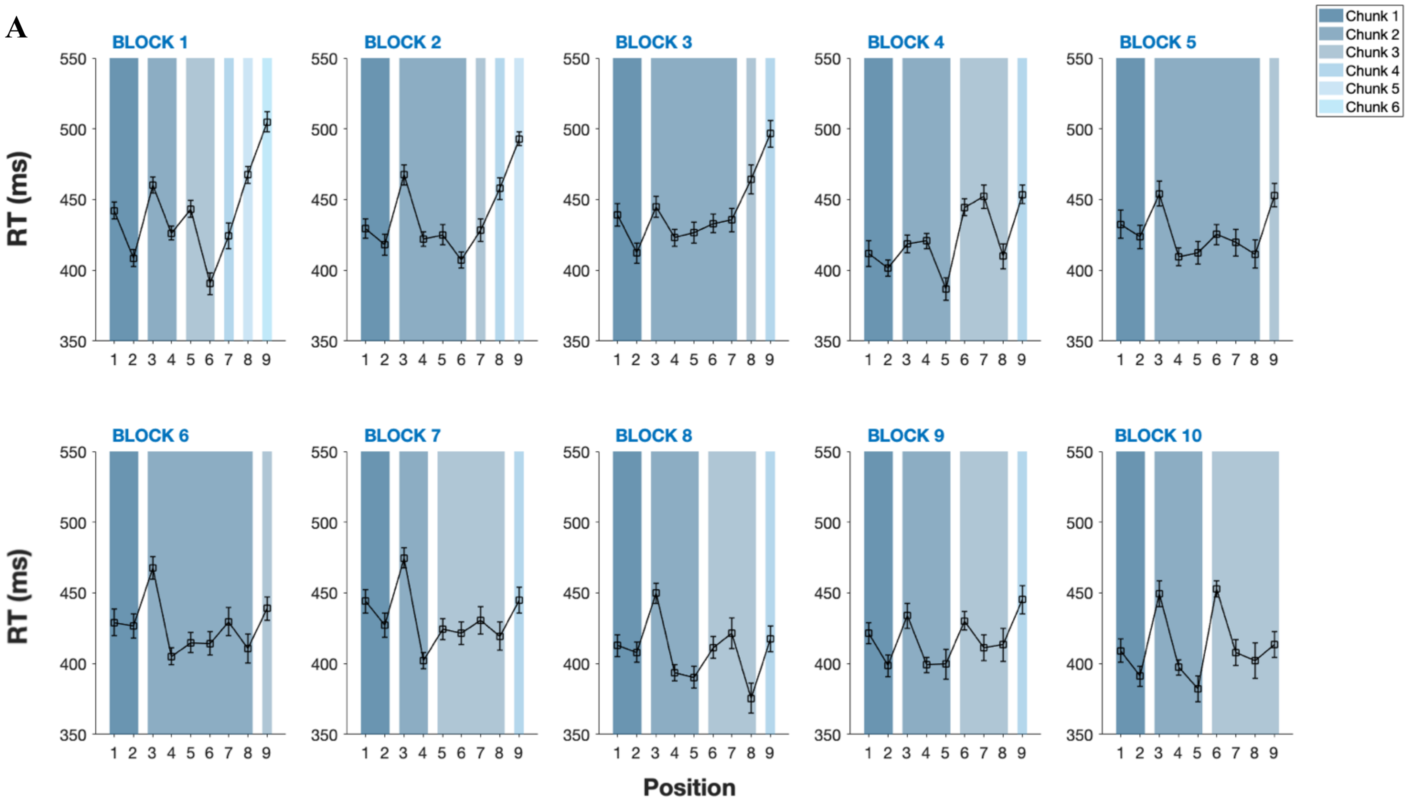


B

| | Chunk size | | | | | | Nb of chunks | Concatenation | Recombination |
|---|---|---|---|---|---|---|---|---|---|
| | C1 | C2 | C3 | C4 | C5 | C6 | | | |
| Block 1 | 2 | 2 | 2 | 1 | 1 | 1 | 6 | - | - |
| 2 | 2 | 4 | 1 | 1 | 1 | - | 5 | 1 | - |
| 3 | 2 | 5 | 1 | 1 | - | - | 4 | 1 | - |
| 4 | 2 | 3 | 3 | 1 | - | - | 4 | 1 | 1 |
| 5 | 2 | 6 | 1 | - | - | - | 3 | 1 | - |
| 6 | 2 | 6 | 1 | - | - | - | 3 | - | - |
| 7 | 2 | 2 | 4 | 1 | - | - | 4 | - | 1 |
| 8 | 2 | 3 | 3 | 1 | - | - | 4 | - | 1 |
| 9 | 2 | 3 | 3 | 1 | - | - | 4 | - | - |
| 10 | 2 | 3 | 4 | - | - | - | 3 | 1 | - |
| Total | | | | | | | | 5 | 3 |

*Note.* A. Mean RT per position across the 10 blocks of trials for one baboon (Angèle) showing the evolution of the chunking pattern (error bars represent 95% confidence intervals). B. Summary table of the reorganizations observed throughout the task.

**Figure and Table S2**

*Evolution of the chunking pattern for Arielle*

**A**

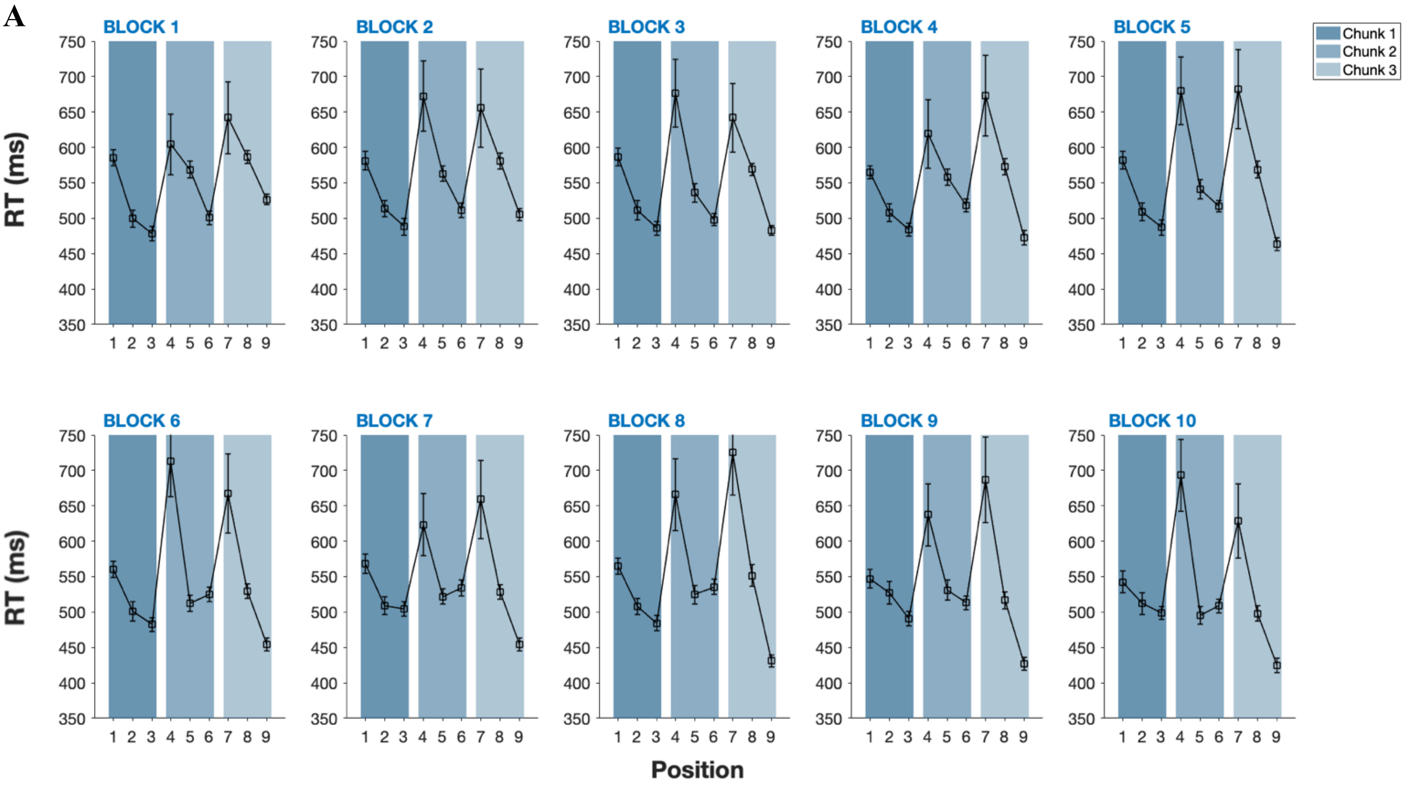


**B**

| | Chunk size | | | Nb of chunks | Concatenation | Recombination |
|---|---|---|---|---|---|---|
| | C1 | C2 | C3 | | | |
| Block 1 | 3 | 3 | 3 | 3 | - | - |
| 2 | 3 | 3 | 3 | 3 | - | - |
| 3 | 3 | 3 | 3 | 3 | - | - |
| 4 | 3 | 3 | 3 | 3 | - | - |
| 5 | 3 | 3 | 3 | 3 | - | - |
| 6 | 3 | 3 | 3 | 3 | - | - |
| 7 | 3 | 3 | 3 | 3 | - | - |
| 8 | 3 | 3 | 3 | 3 | - | - |
| 9 | 3 | 3 | 3 | 3 | - | - |
| 10 | 3 | 3 | 3 | 3 | - | - |
| Total | | | | | 0 | 0 |

*Note.* A. Mean RT per position across the 10 blocks of trials for one baboon (Arielle) showing the evolution of the chunking pattern (error bars represent 95% confidence intervals). B. Summary table of the reorganizations observed throughout the task.

**Figure and Table S3**

*Evolution of the chunking pattern for Cauet*

A

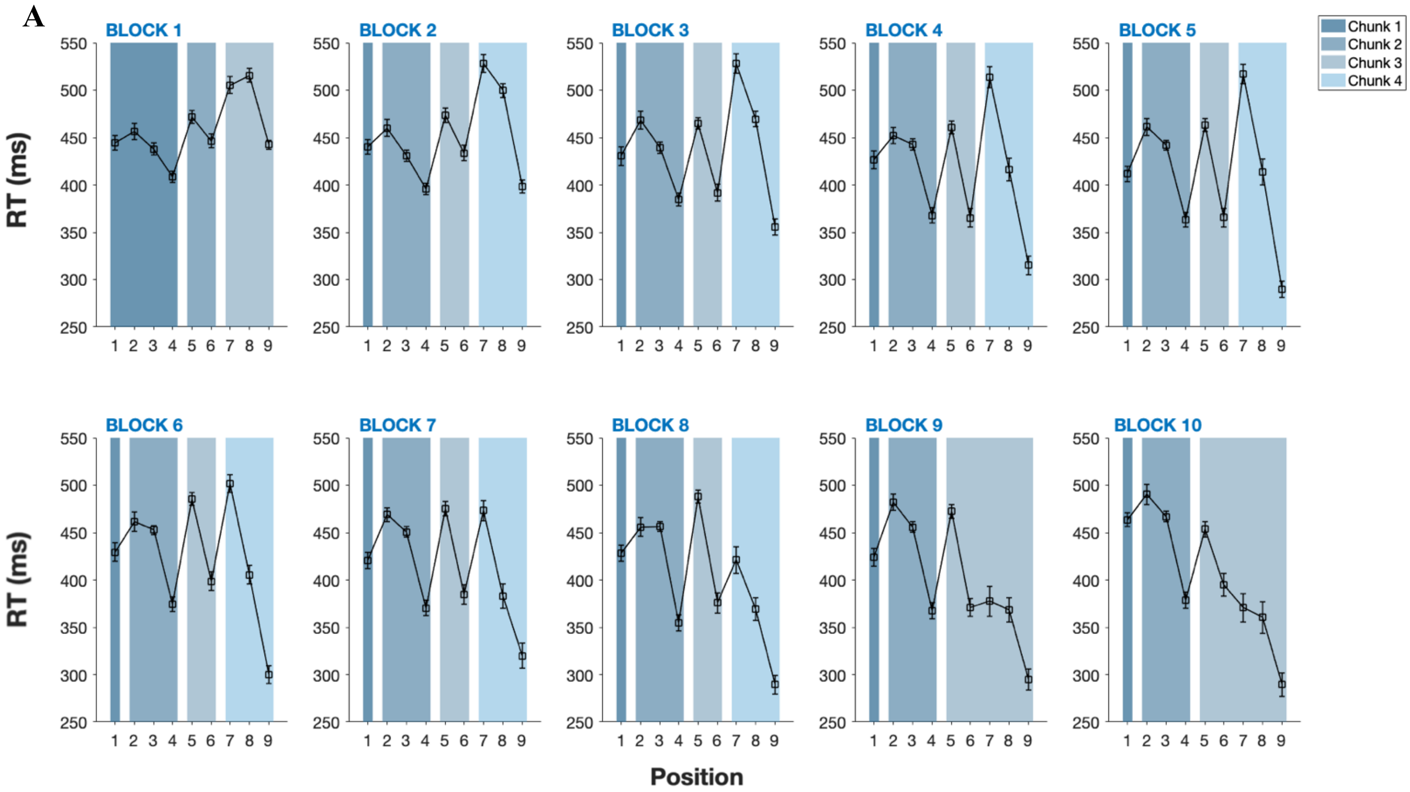


B

| | Chunk size | | | | Nb of chunks | Concatenation | Recombination |
|---|---|---|---|---|---|---|---|
| | C1 | C2 | C3 | C4 | | | |
| Block 1 | 4 | 2 | 3 | - | 3 | - | - |
| 2 | 1 | 3 | 2 | 3 | 4 | - | 1 |
| 3 | 1 | 3 | 2 | 3 | 4 | - | - |
| 4 | 1 | 3 | 2 | 3 | 4 | - | - |
| 5 | 1 | 3 | 2 | 3 | 4 | - | - |
| 6 | 1 | 3 | 2 | 3 | 4 | - | - |
| 7 | 1 | 3 | 2 | 3 | 4 | - | - |
| 8 | 1 | 3 | 2 | 3 | 4 | - | - |
| 9 | 1 | 3 | 5 | - | 3 | 1 | - |
| 10 | 1 | 3 | 5 | - | 3 | - | - |
| Total | | | | | | 1 | 1 |

*Note.* A. Mean RT per position across the 10 blocks of trials for one baboon (Cauet) showing the evolution of the chunking pattern (error bars represent 95% confidence intervals). B. Summary table of the reorganizations observed throughout the task.

**Figure and Table S4**

*Evolution of the chunking pattern for Dream*

**A**

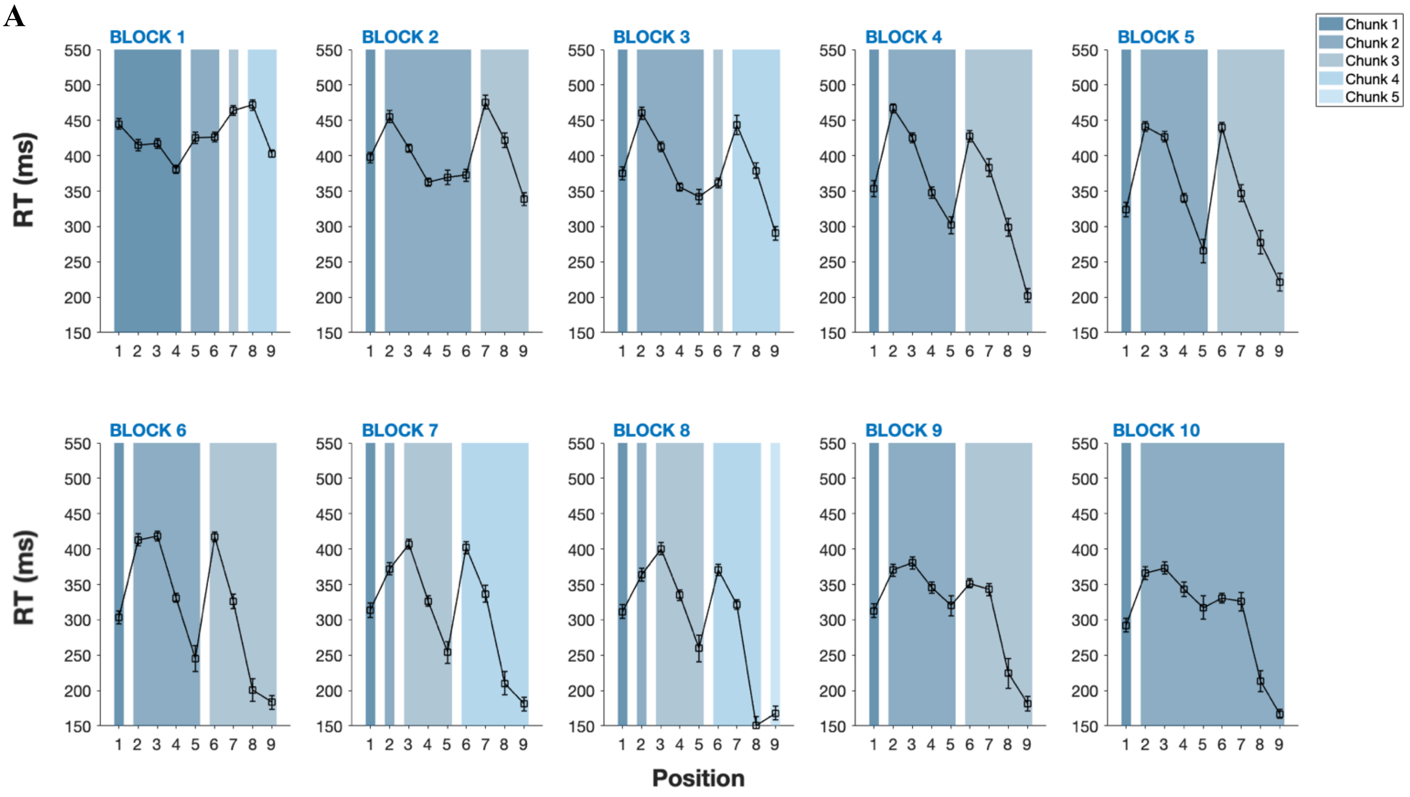


**B**

| | Chunk size | | | | | Nb of chunks | Concatenation | Recombination |
|---|---|---|---|---|---|---|---|---|
| | C1 | C2 | C3 | C4 | C5 | | | |
| Block 1 | 4 | 2 | 1 | 2 | - | 4 | - | - |
| 2 | 1 | 5 | 3 | - | - | 3 | 1 | 1 |
| 3 | 1 | 4 | 1 | 3 | - | 4 | - | 1 |
| 4 | 1 | 4 | 4 | - | - | 3 | 1 | - |
| 5 | 1 | 4 | 4 | - | - | 3 | - | - |
| 6 | 1 | 4 | 4 | - | - | 3 | - | - |
| 7 | 1 | 1 | 3 | 4 | - | 4 | - | 1 |
| 8 | 1 | 1 | 3 | 3 | 1 | 5 | - | 1 |
| 9 | 1 | 4 | 4 | - | - | 3 | 2 | - |
| 10 | 1 | 8 | - | - | - | 2 | 1 | - |
| Total | | | | | | | 5 | 4 |

*Note.* A. Mean RT per position across the 10 blocks of trials for one baboon (Dream) showing the evolution of the chunking pattern (error bars represent 95% confidence intervals). B. Summary table of the reorganizations observed throughout the task.

**Figure and Table S5**

*Evolution of the chunking pattern for Ewine*

**A**

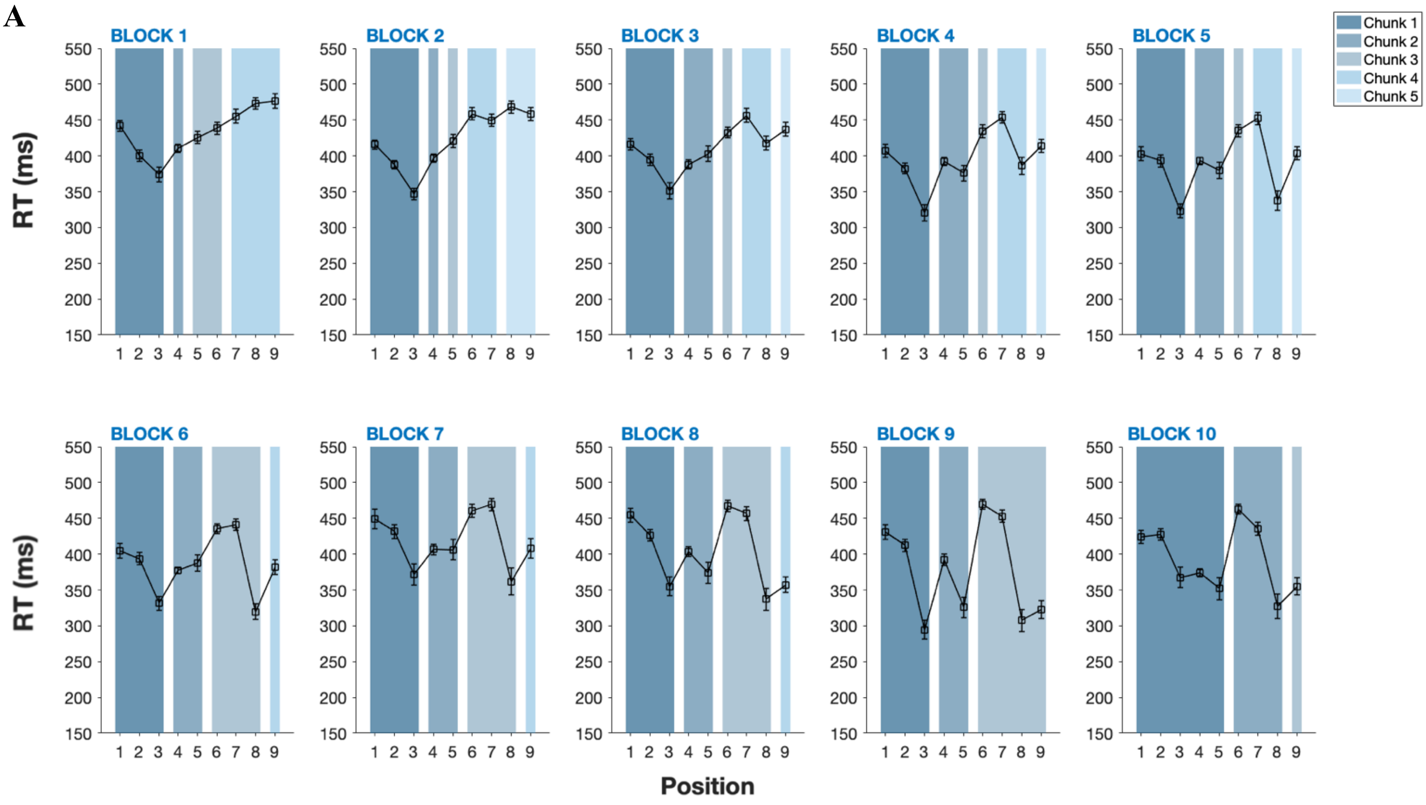


**B**

| | Chunk size | | | | | Nb of chunks | Concatenation | Recombination |
|---|---|---|---|---|---|---|---|---|
| | C1 | C2 | C3 | C4 | C5 | | | |
| Block 1 | 3 | 1 | 2 | 3 | - | 4 | - | - |
| 2 | 3 | 1 | 1 | 2 | 2 | 5 | - | 2 |
| 3 | 3 | 2 | 1 | 2 | 1 | 5 | 1 | 1 |
| 4 | 3 | 2 | 1 | 2 | 1 | 5 | - | - |
| 5 | 3 | 2 | 1 | 2 | 1 | 5 | - | - |
| 6 | 3 | 2 | 3 | 1 | - | 4 | 1 | - |
| 7 | 3 | 2 | 3 | 1 | - | 4 | - | - |
| 8 | 3 | 2 | 3 | 1 | - | 4 | - | - |
| 9 | 3 | 2 | 4 | - | - | 3 | 1 | - |
| 10 | 5 | 3 | 1 | - | - | 3 | 1 | 1 |
| Total | | | | | | | 4 | 4 |

*Note.* A. Mean RT per position across the 10 blocks of trials for one baboon (Ewine) showing the evolution of the chunking pattern (error bars represent 95% confidence intervals). B. Summary table of the reorganizations observed throughout the task.

### Figure and Table S6

*Evolution of the chunking pattern for Fana*

A

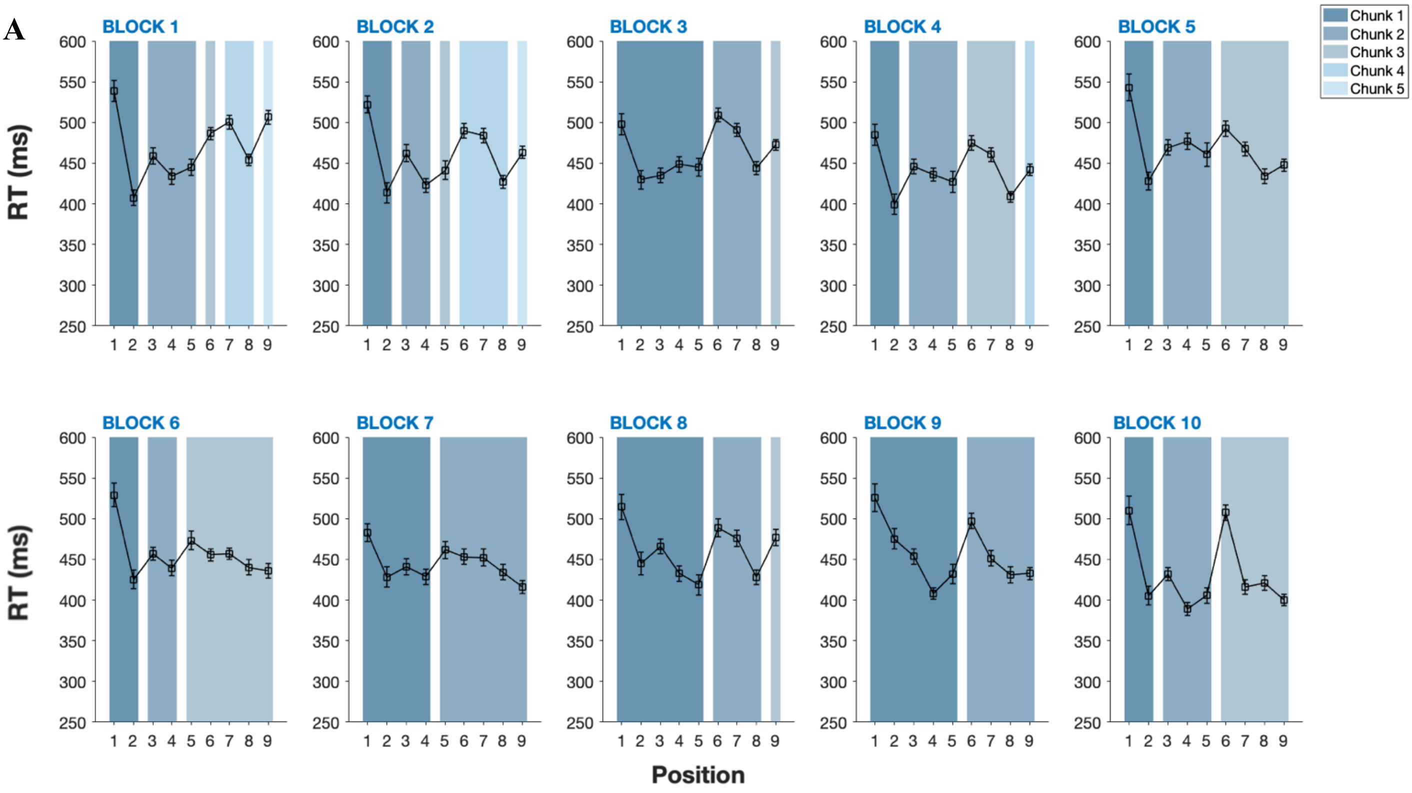


B

| | Chunk size | | | | | Nb of chunks | Concatenation | Recombination |
|---|---|---|---|---|---|---|---|---|
| | C1 | C2 | C3 | C4 | C5 | | | |
| Block 1 | 2 | 3 | 1 | 2 | 1 | 5 | - | - |
| 2 | 2 | 2 | 1 | 3 | 1 | 5 | 1 | 1 |
| 3 | 5 | 3 | 1 | - | - | 3 | 2 | - |
| 4 | 2 | 3 | 3 | 1 | - | 4 | - | 1 |
| 5 | 2 | 3 | 4 | - | - | 3 | 1 | - |
| 6 | 2 | 2 | 5 | - | - | 3 | - | 1 |
| 7 | 4 | 5 | - | - | - | 2 | 1 | - |
| 8 | 5 | 3 | 1 | - | - | 3 | - | 2 |
| 9 | 5 | 4 | - | - | - | 2 | 1 | - |
| 10 | 2 | 3 | 4 | - | - | 3 | - | 1 |
| Total | | | | | | | 6 | 6 |

*Note.* A. Mean RT per position across the 10 blocks of trials for one baboon (Fana) showing the evolution of the chunking pattern (error bars represent 95% confidence intervals). B. Summary table of the reorganizations observed throughout the task.

**Figure and Table S7**

*Evolution of the chunking pattern for Felipe*

A

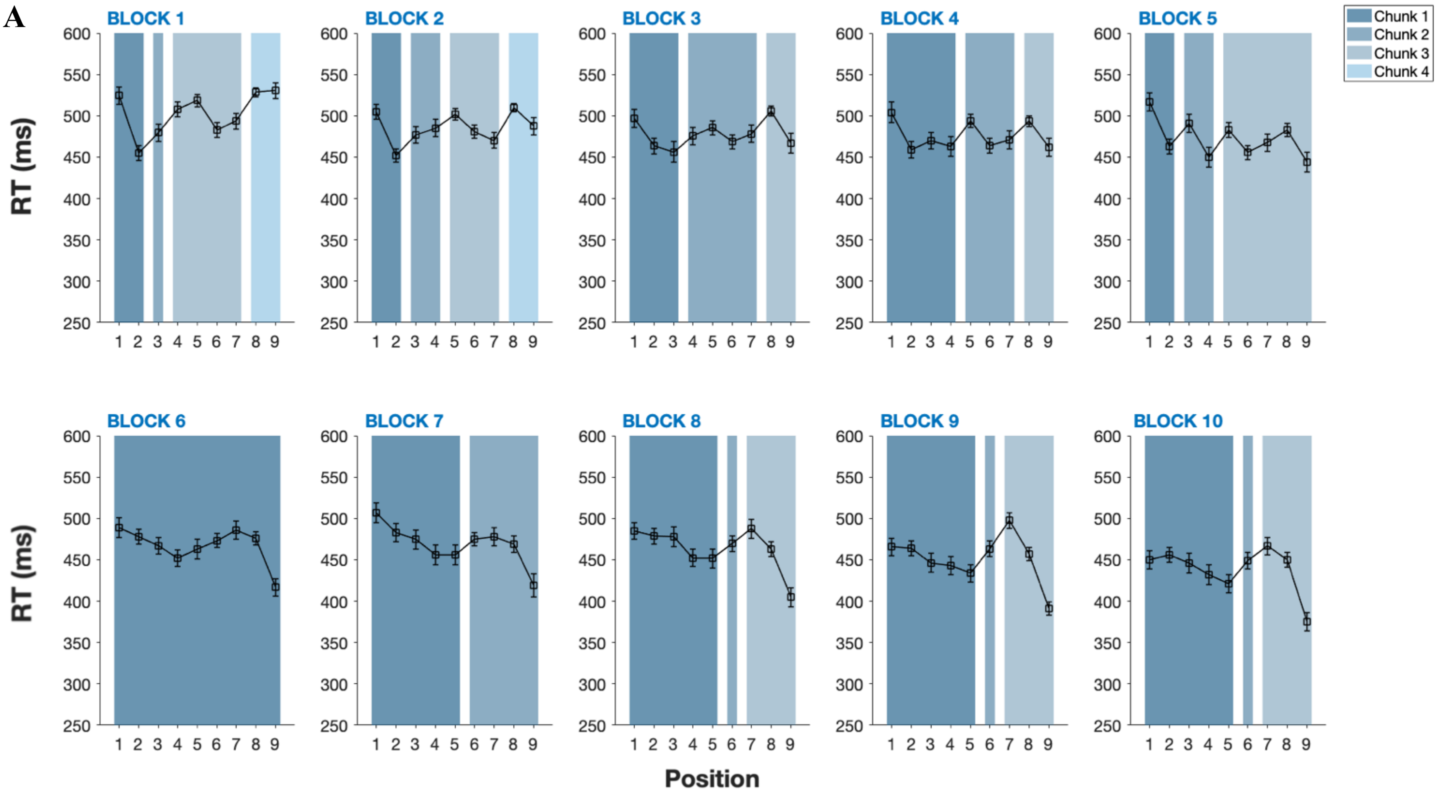


B

| | Chunk size | | | | Nb of chunks | Concatenation | Recombination |
|---|---|---|---|---|---|---|---|
| | C1 | C2 | C3 | C4 | | | |
| Block 1 | 2 | 1 | 4 | 2 | 4 | - | - |
| 2 | 2 | 2 | 3 | 2 | 4 | - | 1 |
| 3 | 3 | 4 | 2 | - | 3 | - | 1 |
| 4 | 4 | 3 | 2 | - | 3 | - | 1 |
| 5 | 2 | 2 | 5 | - | 3 | 1 | 1 |
| 6 | 9 | - | - | - | 1 | 2 | - |
| 7 | 5 | 4 | - | - | 2 | - | 1 |
| 8 | 5 | 1 | 3 | - | 3 | - | 1 |
| 9 | 5 | 1 | 3 | - | 3 | - | - |
| 10 | 5 | 1 | 3 | - | 3 | - | - |
| Total | | | | | | 3 | 6 |

*Note.* A. Mean RT per position across the 10 blocks of trials for one baboon (Felipe) showing the evolution of the chunking pattern (error bars represent 95% confidence intervals). B. Summary table of the reorganizations observed throughout the task.

**Figure and Table S8**

*Evolution of the chunking pattern for Feya*

A

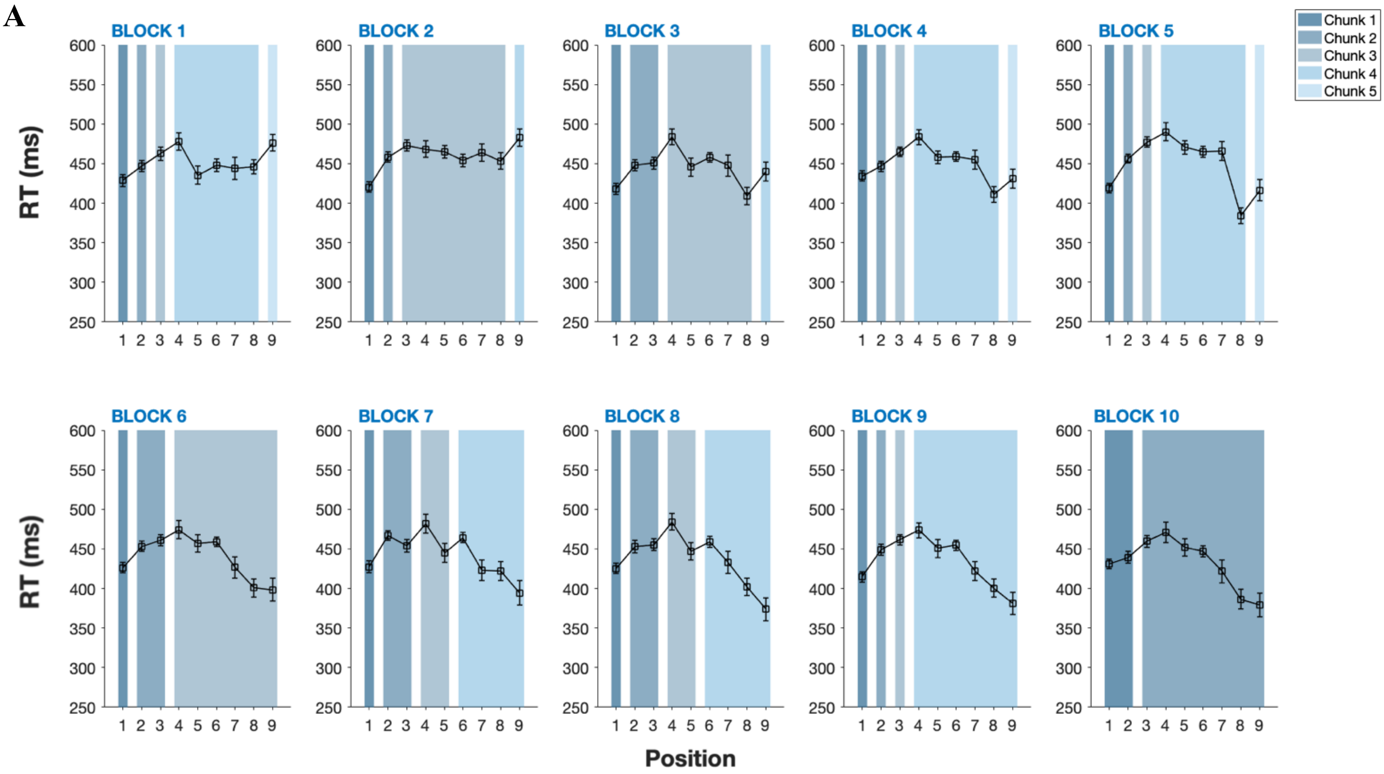


B

| | Chunk size | | | | | Nb of chunks | Concatenation | Recombination |
|---|---|---|---|---|---|---|---|---|
| | C1 | C2 | C3 | C4 | C5 | | | |
| Block 1 | 1 | 1 | 1 | 5 | 1 | 5 | - | - |
| 2 | 1 | 1 | 6 | 1 | - | 4 | 1 | - |
| 3 | 1 | 2 | 5 | 1 | - | 4 | - | 1 |
| 4 | 1 | 1 | 1 | 5 | 1 | 5 | - | 1 |
| 5 | 1 | 1 | 1 | 5 | 1 | 5 | - | - |
| 6 | 1 | 2 | 6 | - | - | 3 | 2 | - |
| 7 | 1 | 2 | 2 | 4 | - | 4 | - | 1 |
| 8 | 1 | 2 | 2 | 4 | - | 4 | - | - |
| 9 | 1 | 1 | 1 | 6 | - | 4 | - | 2 |
| 10 | 2 | 7 | - | - | - | 2 | 1 | 1 |
| Total | | | | | | | 4 | 6 |

*Note.* A. Mean RT per position across the 10 blocks of trials for one baboon (Feya) showing the evolution of the chunking pattern (error bars represent 95% confidence intervals). B. Summary table of the reorganizations observed throughout the task.

**Figure and Table S9**

*Evolution of the chunking pattern for Flute*

A

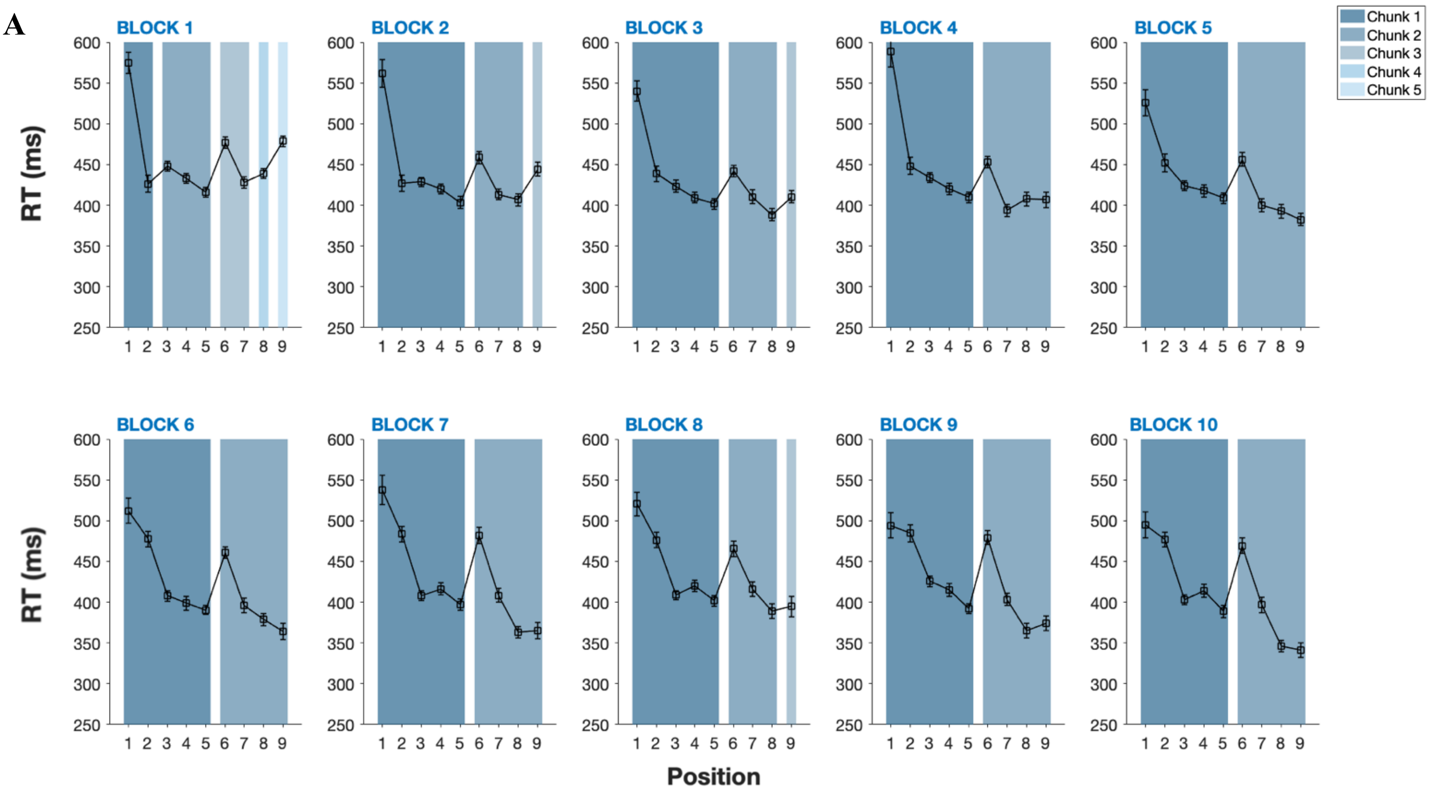


B

| | Chunk size | | | | | Nb of chunks | Concatenation | Recombination |
|---|---|---|---|---|---|---|---|---|
| | C1 | C2 | C3 | C4 | C5 | | | |
| Block 1 | 2 | 3 | 2 | 1 | 1 | 5 | - | - |
| 2 | 5 | 3 | 1 | - | - | 3 | 2 | - |
| 3 | 5 | 3 | 1 | - | - | 3 | - | - |
| 4 | 5 | 4 | - | - | - | 2 | 1 | - |
| 5 | 5 | 4 | - | - | - | 2 | - | - |
| 6 | 5 | 4 | - | - | - | 2 | - | - |
| 7 | 5 | 4 | - | - | - | 2 | - | - |
| 8 | 5 | 3 | 1 | - | - | 3 | - | 1 |
| 9 | 5 | 4 | - | - | - | 2 | 1 | - |
| 10 | 5 | 4 | - | - | - | 2 | - | - |
| Total | | | | | | | 4 | 1 |

*Note.* A. Mean RT per position across the 10 blocks of trials for one baboon (Flute) showing the evolution of the chunking pattern (error bars represent 95% confidence intervals). B. Summary table of the reorganizations observed throughout the task.

**Figure and Table S10**

*Evolution of the chunking pattern for Harlem*

**A**

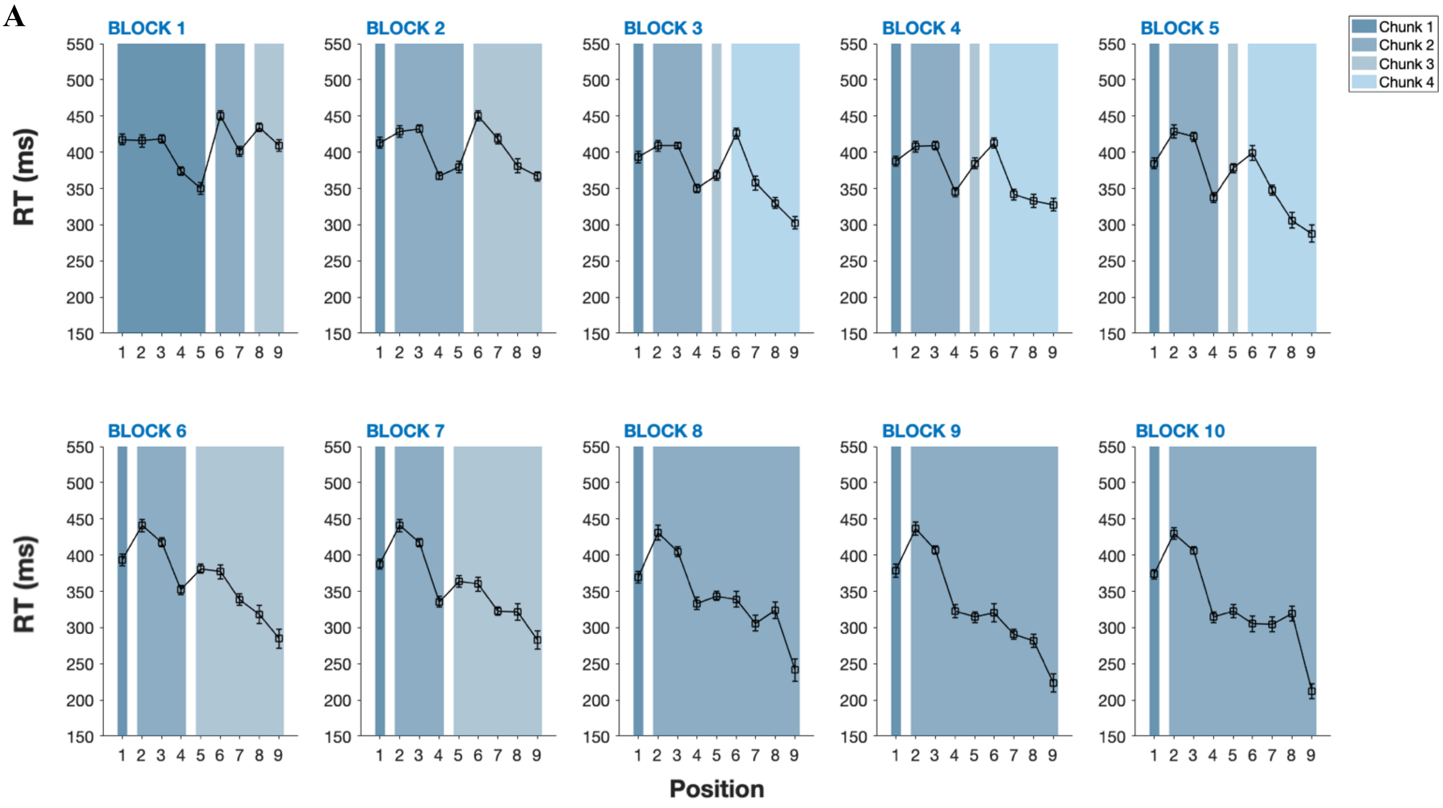


**B**

| | Chunk size | | | | Nb of chunks | Concatenation | Recombination |
|---|---|---|---|---|---|---|---|
| | C1 | C2 | C3 | C4 | | | |
| Block 1 | 5 | 2 | 2 | - | 3 | - | - |
| 2 | 1 | 4 | 4 | - | 3 | 1 | 1 |
| 3 | 1 | 3 | 1 | 4 | 4 | - | 1 |
| 4 | 1 | 3 | 1 | 4 | 4 | - | - |
| 5 | 1 | 3 | 1 | 4 | 4 | - | - |
| 6 | 1 | 3 | 5 | - | 3 | 1 | - |
| 7 | 1 | 3 | 5 | - | 3 | - | - |
| 8 | 1 | 8 | - | - | 2 | 1 | - |
| 9 | 1 | 8 | - | - | 2 | - | - |
| 10 | 1 | 8 | - | - | 2 | - | - |
| Total | | | | | | 3 | 2 |

*Note.* A. Mean RT per position across the 10 blocks of trials for one baboon (Harlem) showing the evolution of the chunking pattern (error bars represent 95% confidence intervals). B. Summary table of the reorganizations observed throughout the task.

### Figure and Table S11

*Evolution of the chunking pattern for Kali*

A

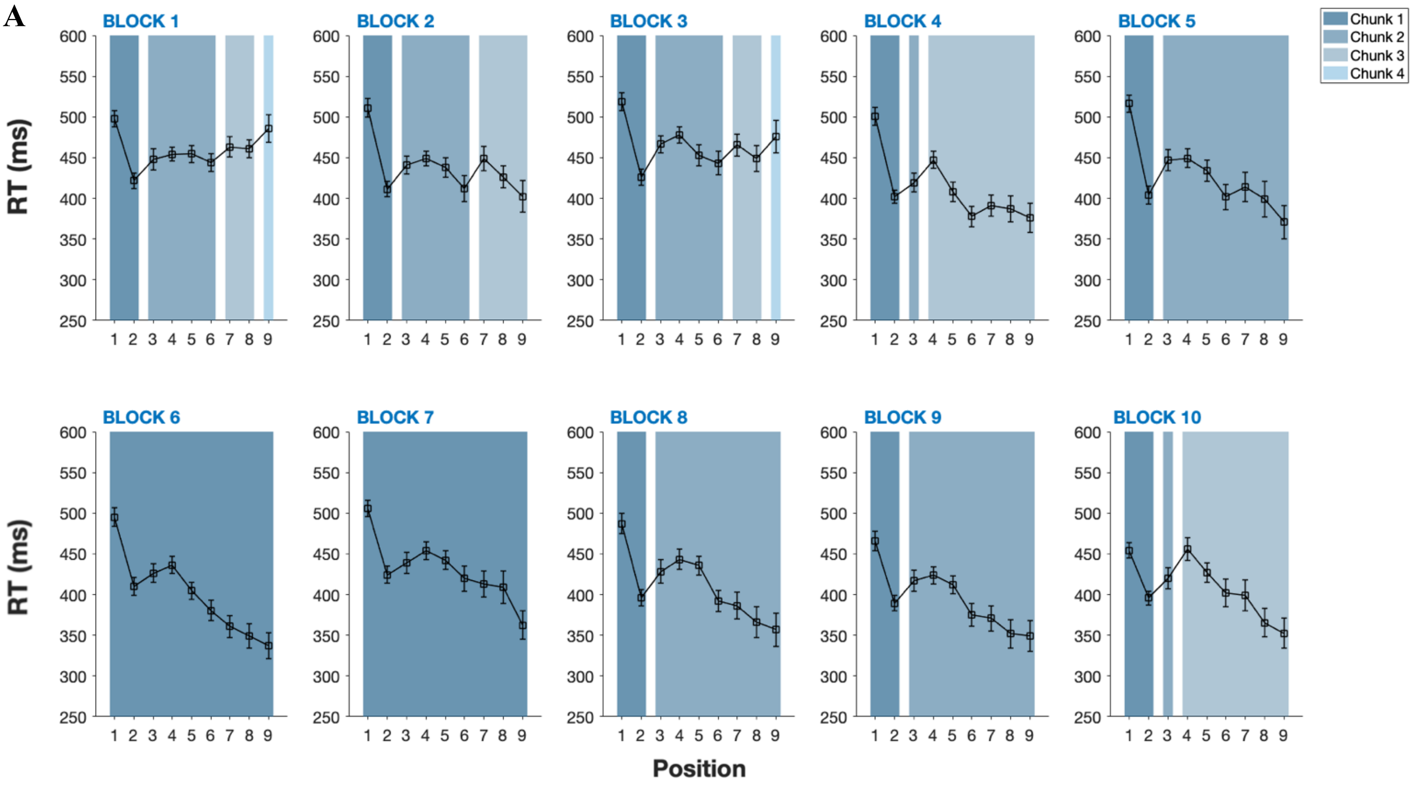


B

| | Chunk size | | | | Nb of chunks | Concatenation | Recombination |
|---|---|---|---|---|---|---|---|
| | C1 | C2 | C3 | C4 | | | |
| Block 1 | 2 | 4 | 2 | 1 | 4 | - | - |
| 2 | 2 | 4 | 3 | - | 3 | 1 | - |
| 3 | 2 | 4 | 2 | 1 | 4 | - | 1 |
| 4 | 2 | 1 | 6 | - | 3 | 1 | 1 |
| 5 | 2 | 7 | - | - | 2 | 1 | - |
| 6 | 9 | - | - | - | 1 | 1 | - |
| 7 | 9 | - | - | - | 1 | - | - |
| 8 | 2 | 7 | - | - | 2 | - | 1 |
| 9 | 2 | 7 | - | - | 2 | - | - |
| 10 | 2 | 1 | 6 | - | 3 | - | 1 |
| Total | | | | | | 4 | 4 |

*Note.* A. Mean RT per position across the 10 blocks of trials for one baboon (Kali) showing the evolution of the chunking pattern (error bars represent 95% confidence intervals). B. Summary table of the reorganizations observed throughout the task.

**Figure and Table S12**

*Evolution of the chunking pattern for Lips*

A

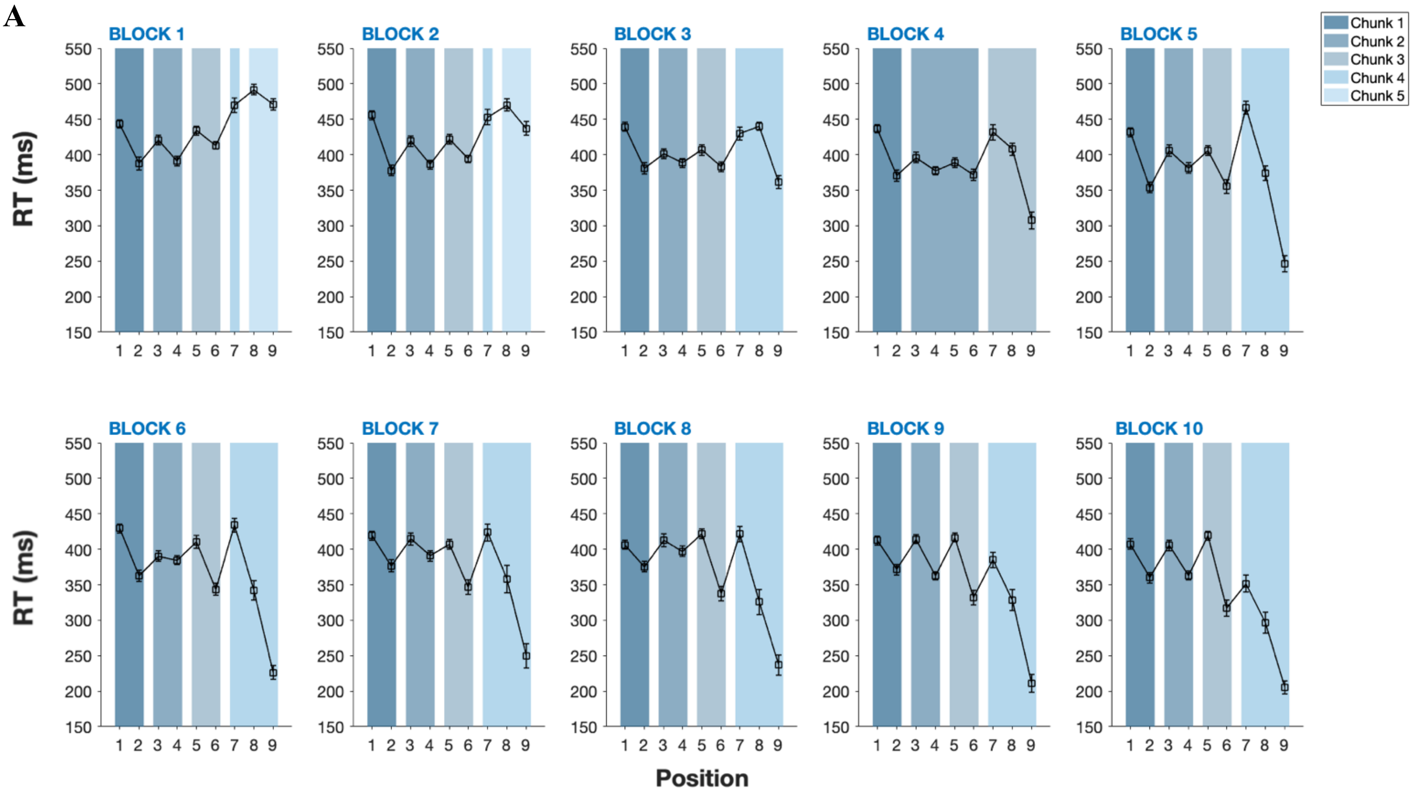


B

| | Chunk size | | | | | Nb of chunks | Concatenation | Recombination |
|---|---|---|---|---|---|---|---|---|
| | C1 | C2 | C3 | C4 | C5 | | | |
| Block 1 | 2 | 2 | 2 | 2 | 1 | 5 | - | - |
| 2 | 2 | 2 | 2 | 2 | - | 5 | - | - |
| 3 | 2 | 2 | 2 | - | - | 4 | 1 | - |
| 4 | 2 | 2 | 2 | - | 1 | 4 | - | - |
| 5 | 2 | 2 | 2 | - | 1 | 4 | - | - |
| 6 | 2 | 2 | 2 | - | - | 4 | - | - |
| 7 | 2 | 2 | 2 | - | - | 4 | - | - |
| 8 | 2 | 2 | 2 | - | - | 4 | - | - |
| 9 | 2 | 2 | 2 | - | - | 4 | - | - |
| 10 | 2 | 2 | 2 | - | - | 4 | - | - |
| Total | | | | | | | 1 | 0 |

*Note.* A. Mean RT per position across the 10 blocks of trials for one baboon (Lips) showing the evolution of the chunking pattern (error bars represent 95% confidence intervals). B. Summary table of the reorganizations observed throughout the task.

**Figure and Table S13**

*Evolution of the chunking pattern for Lome*

**A**

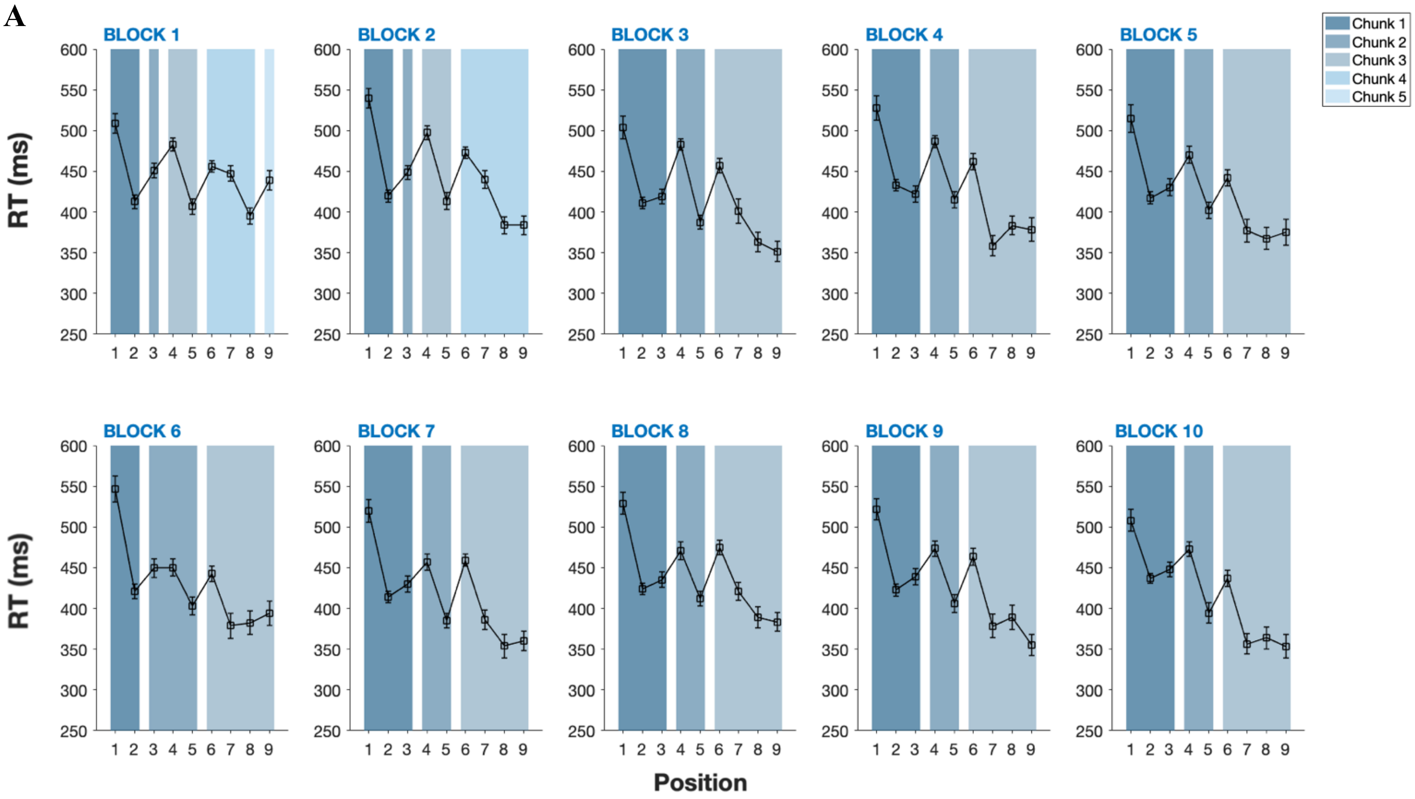


**B**

| | Chunk size | | | | | Nb of chunks | Concatenation | Recombination |
|---|---|---|---|---|---|---|---|---|
| | C1 | C2 | C3 | C4 | C5 | | | |
| Block 1 | 2 | 1 | 2 | 3 | 1 | 5 | - | - |
| 2 | 2 | 1 | 2 | 4 | - | 4 | 1 | - |
| 3 | 3 | 2 | 4 | - | - | 3 | 1 | - |
| 4 | 3 | 2 | 4 | - | - | 3 | - | - |
| 5 | 3 | 2 | 4 | - | - | 3 | - | - |
| 6 | 3 | 2 | 4 | - | - | 3 | - | - |
| 7 | 3 | 2 | 4 | - | - | 3 | - | - |
| 8 | 3 | 2 | 4 | - | - | 3 | - | - |
| 9 | 3 | 2 | 4 | - | - | 3 | - | - |
| 10 | 3 | 2 | 4 | - | - | 3 | - | - |
| Total | | | | | | | 2 | 0 |

*Note.* A. Mean RT per position across the 10 blocks of trials for one baboon (Lome) showing the evolution of the chunking pattern (error bars represent 95% confidence intervals). B. Summary table of the reorganizations observed throughout the task.

**Figure and Table S14**

*Evolution of the chunking pattern for Mako*

**A**

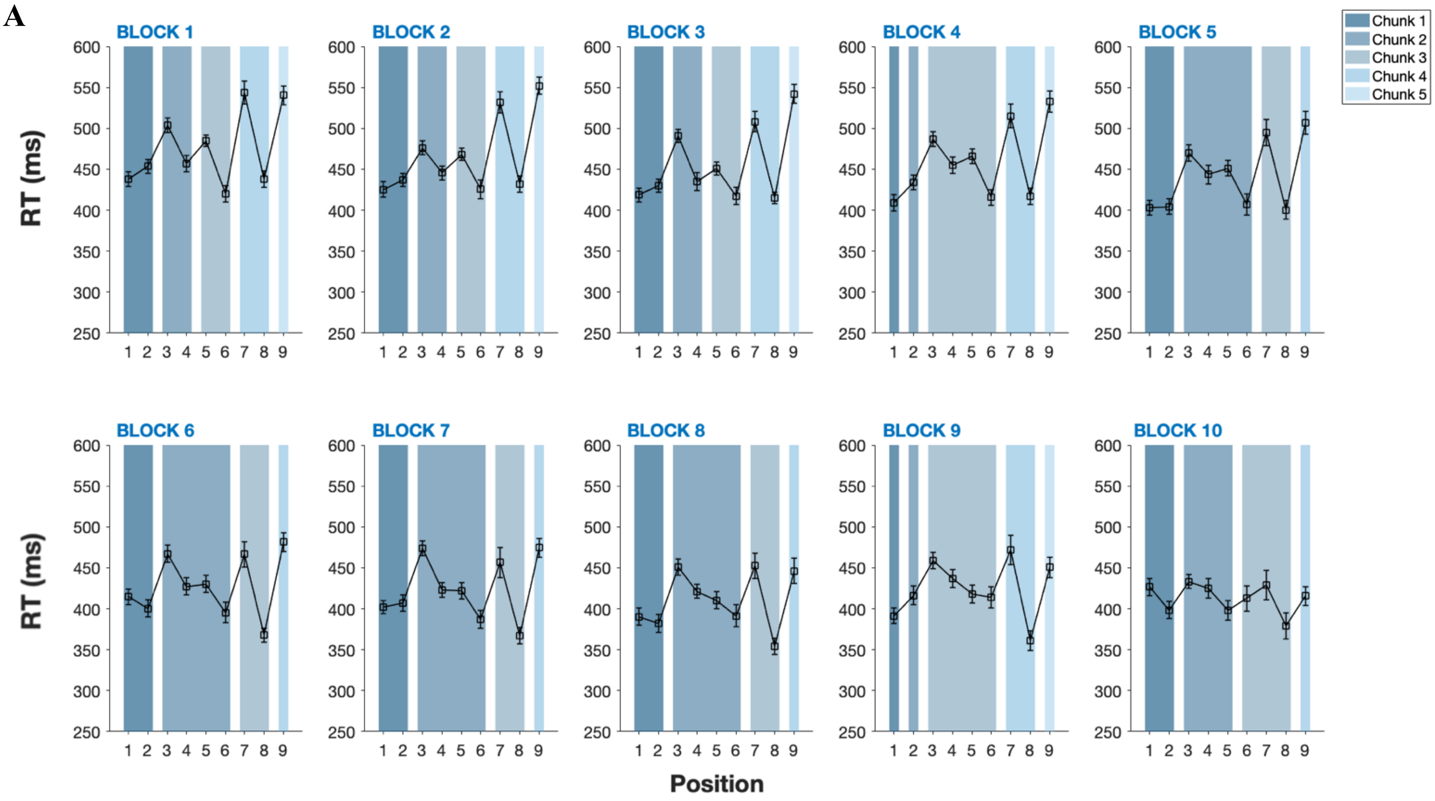


**B**

| | Chunk size | | | | | Nb of chunks | Concatenation | Recombination |
|---|---|---|---|---|---|---|---|---|
| | C1 | C2 | C3 | C4 | C5 | | | |
| Block 1 | 2 | 2 | 2 | 2 | 1 | 5 | - | - |
| 2 | 2 | 2 | 2 | 2 | 1 | 5 | - | - |
| 3 | 2 | 2 | 2 | 2 | 1 | 5 | - | - |
| 4 | 1 | 1 | 4 | 2 | 1 | 5 | 1 | 1 |
| 5 | 2 | 4 | 2 | 1 | - | 4 | 1 | - |
| 6 | 2 | 4 | 2 | 1 | - | 4 | - | - |
| 7 | 2 | 4 | 2 | 1 | - | 4 | - | - |
| 8 | 2 | 4 | 2 | 1 | - | 4 | - | - |
| 9 | 1 | 1 | 4 | 2 | 1 | 5 | - | 1 |
| 10 | 2 | 3 | 3 | 1 | - | 4 | 1 | 1 |
| Total | | | | | | | 3 | 3 |

*Note.* A. Mean RT per position across the 10 blocks of trials for one baboon (Mako) showing the evolution of the chunking pattern (error bars represent 95% confidence intervals). B. Summary table of the reorganizations observed throughout the task.

**Figure and Table S15**

*Evolution of the chunking pattern for Mali*

A

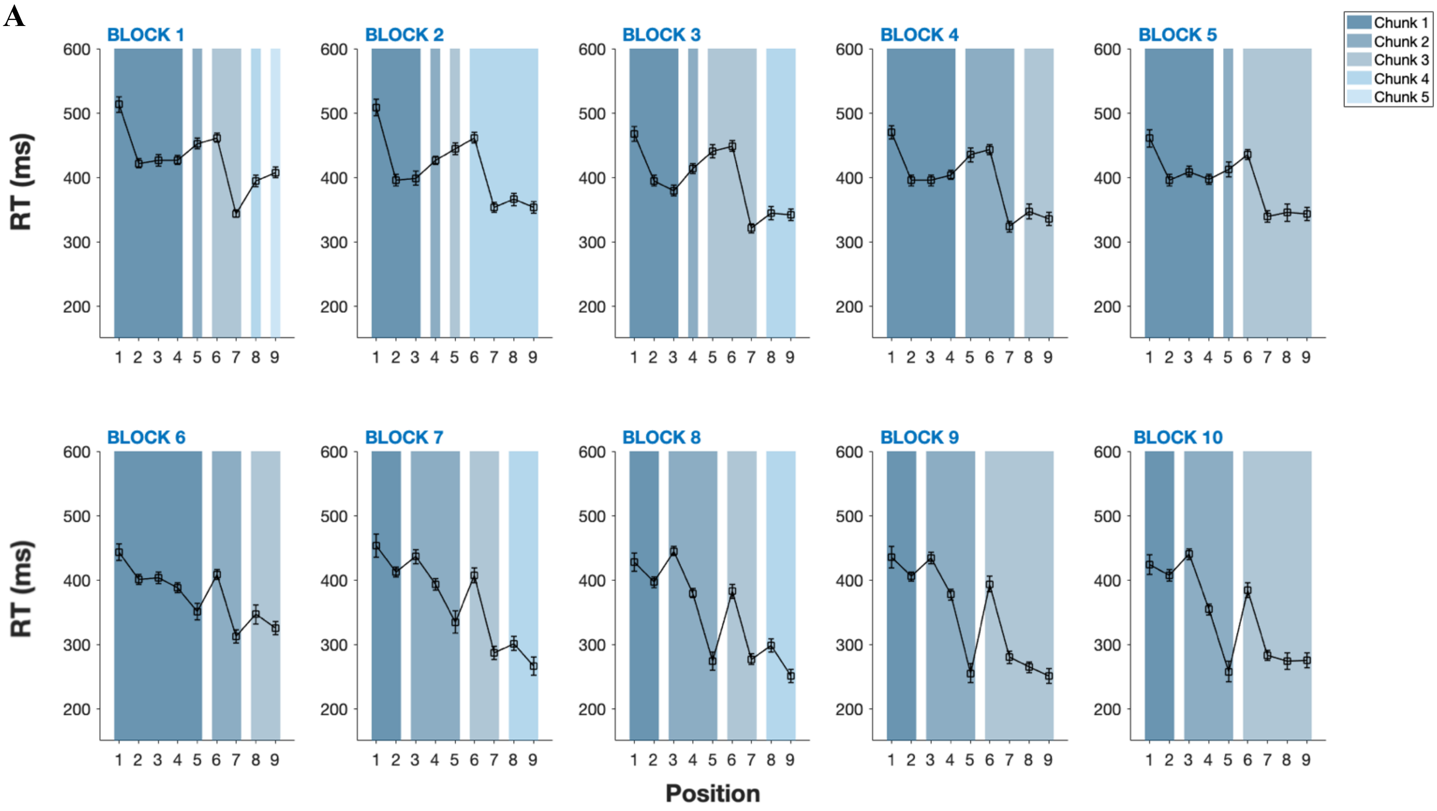


B

| | Chunk size | | | | | Nb of chunks | Concatenation | Recombination |
|---|---|---|---|---|---|---|---|---|
| | C1 | C2 | C3 | C4 | C5 | | | |
| Block 1 | 4 | 1 | 2 | 1 | 1 | 5 | - | - |
| 2 | 3 | 1 | 1 | 4 | - | 4 | 2 | 1 |
| 3 | 3 | 1 | 3 | 2 | - | 4 | 1 | 1 |
| 4 | 4 | 3 | 2 | - | - | 3 | 1 | - |
| 5 | 4 | 1 | 4 | - | - | 3 | - | 2 |
| 6 | 5 | 2 | 2 | - | - | 3 | 1 | 1 |
| 7 | 2 | 3 | 2 | 2 | - | 4 | - | 1 |
| 8 | 2 | 3 | 2 | 2 | - | 4 | - | - |
| 9 | 2 | 3 | 4 | - | - | 3 | 1 | - |
| 10 | 2 | 3 | 4 | - | - | 3 | - | - |
| Total | | | | | | | 6 | 6 |

*Note.* A. Mean RT per position across the 10 blocks of trials for one baboon (Mali) showing the evolution of the chunking pattern (error bars represent 95% confidence intervals). B. Summary table of the reorganizations observed throughout the task.

**Figure and Table S16**

*Evolution of the chunking pattern for Muse*

A

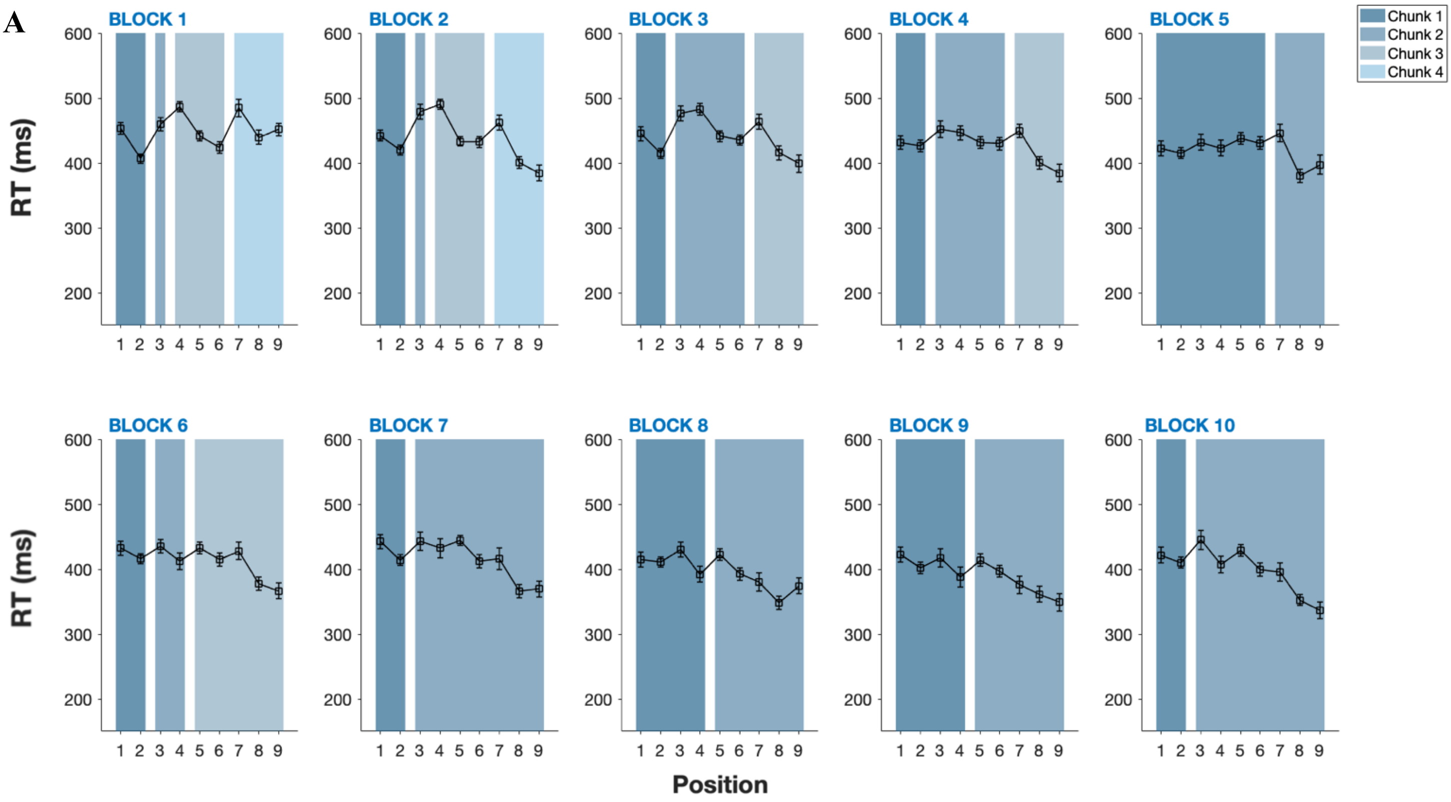


B

| | Chunk size | | | | Nb of chunks | Concatenation | Recombination |
|---|---|---|---|---|---|---|---|
| | C1 | C2 | C3 | C4 | | | |
| Block 1 | 2 | 1 | 3 | 3 | 4 | - | - |
| 2 | 2 | 1 | 3 | 3 | 4 | - | - |
| 3 | 2 | 4 | 3 | - | 3 | 1 | - |
| 4 | 2 | 4 | 3 | - | 3 | - | - |
| 5 | 6 | 3 | - | - | 2 | 1 | - |
| 6 | 2 | 2 | 5 | - | 3 | - | 3 |
| 7 | 2 | 7 | - | - | 2 | 1 | - |
| 8 | 4 | 5 | - | - | 2 | - | 1 |
| 9 | 4 | 5 | - | - | 2 | - | - |
| 10 | 2 | 7 | - | - | 2 | - | 1 |
| Total | | | | | | 3 | 5 |

*Note.* A. Mean RT per position across the 10 blocks of trials for one baboon (Muse) showing the evolution of the chunking pattern (error bars represent 95% confidence intervals). B. Summary table of the reorganizations observed throughout the task.

#### Figure and Table S17

*Evolution of the chunking pattern for Violette*

**A**

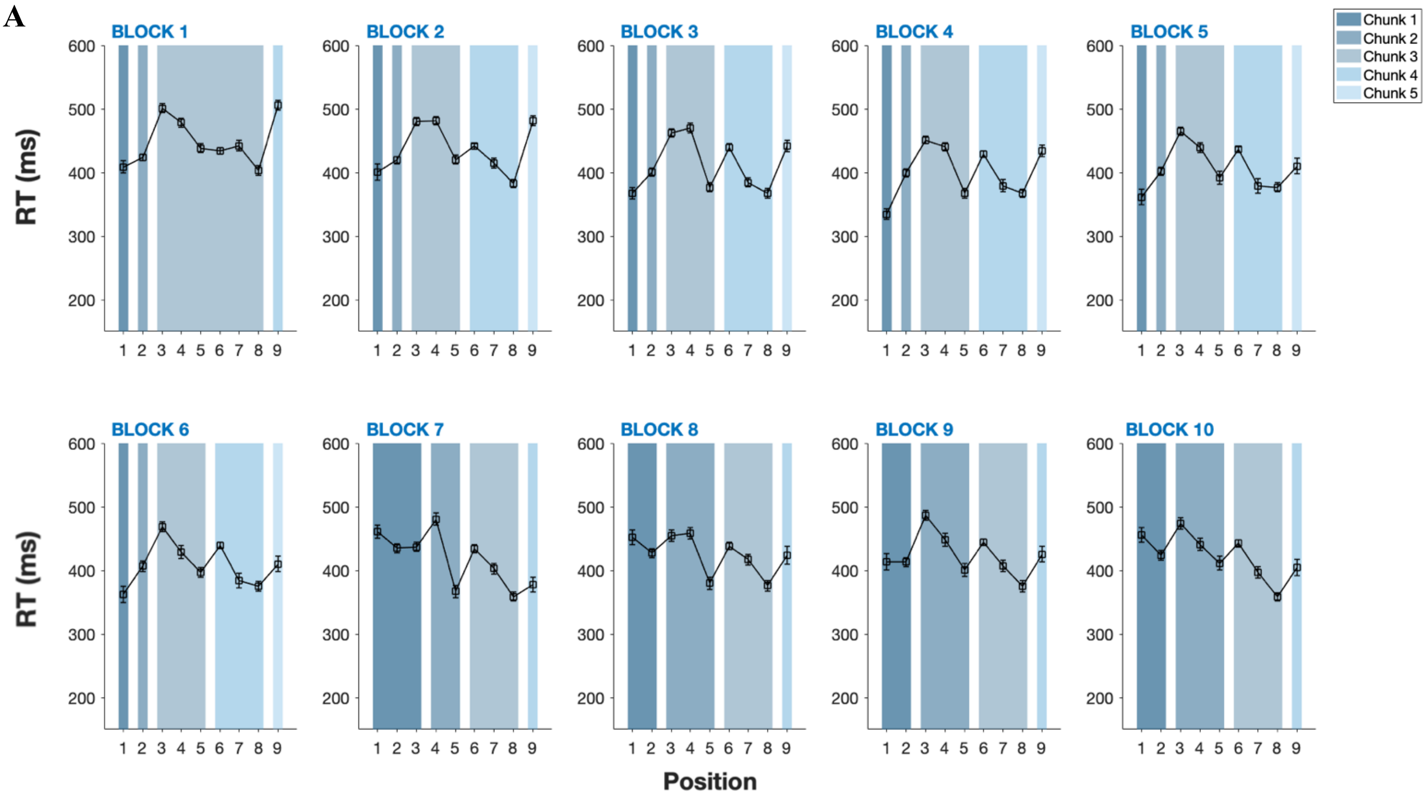


**B**

| | Chunk size | | | | | Nb of chunks | Concatenation | Recombination |
|---|---|---|---|---|---|---|---|---|
| | C1 | C2 | C3 | C4 | C5 | | | |
| Block 1 | 1 | 1 | 6 | 1 | | 4 | - | - |
| 2 | 1 | 1 | 3 | 3 | 1 | 5 | - | 1 |
| 3 | 1 | 1 | 3 | 3 | 1 | 5 | - | - |
| 4 | 1 | 1 | 3 | 3 | 1 | 5 | - | - |
| 5 | 1 | 1 | 3 | 3 | 1 | 5 | - | - |
| 6 | 1 | 1 | 3 | 3 | 1 | 5 | - | - |
| 7 | 3 | 2 | 3 | 1 | - | 4 | 1 | 1 |
| 8 | 2 | 3 | 3 | 1 | - | 4 | - | 1 |
| 9 | 2 | 3 | 3 | 1 | - | 4 | - | - |
| 10 | 2 | 3 | 3 | 1 | - | 4 | - | - |
| Total | | | | | | | 1 | 3 |

*Note.* A. Mean RT per position across the 10 blocks of trials for one baboon (Violette) showing the evolution of the chunking pattern (error bars represent 95% confidence intervals). B. Summary table of the reorganizations observed throughout the task.

**Figure S18**

*Expected transition times (TTs) computed from the random trials and TTs observed at the beginning of the task for the repeated sequences.*

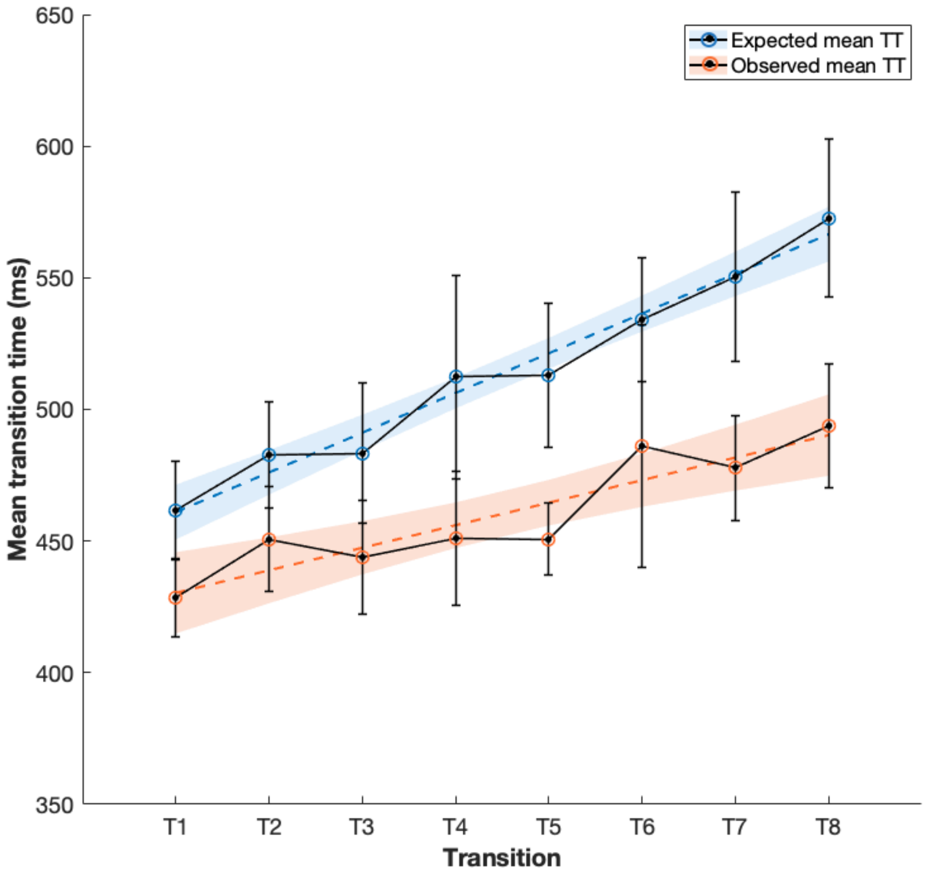


*Note*. The upper blue curve represents the expected mean response time (averaged over the two sequences) computed over the 1,000 initial random trials for each of the eight possible transitions (see Method Section, "Design of the sequences"). The lower orange curve represents the mean response time for each transition that the baboons produced during the first 10 repetitions of the sequence (averaged over the two sequences). This ensures that our sequences were correctly designed and that the ascending pattern of response times was replicated at the start of the experiment. Error bars represent 95% confidence intervals.

**Figure S19**

*Mean RTs for the random and repeated sequences over blocks.*

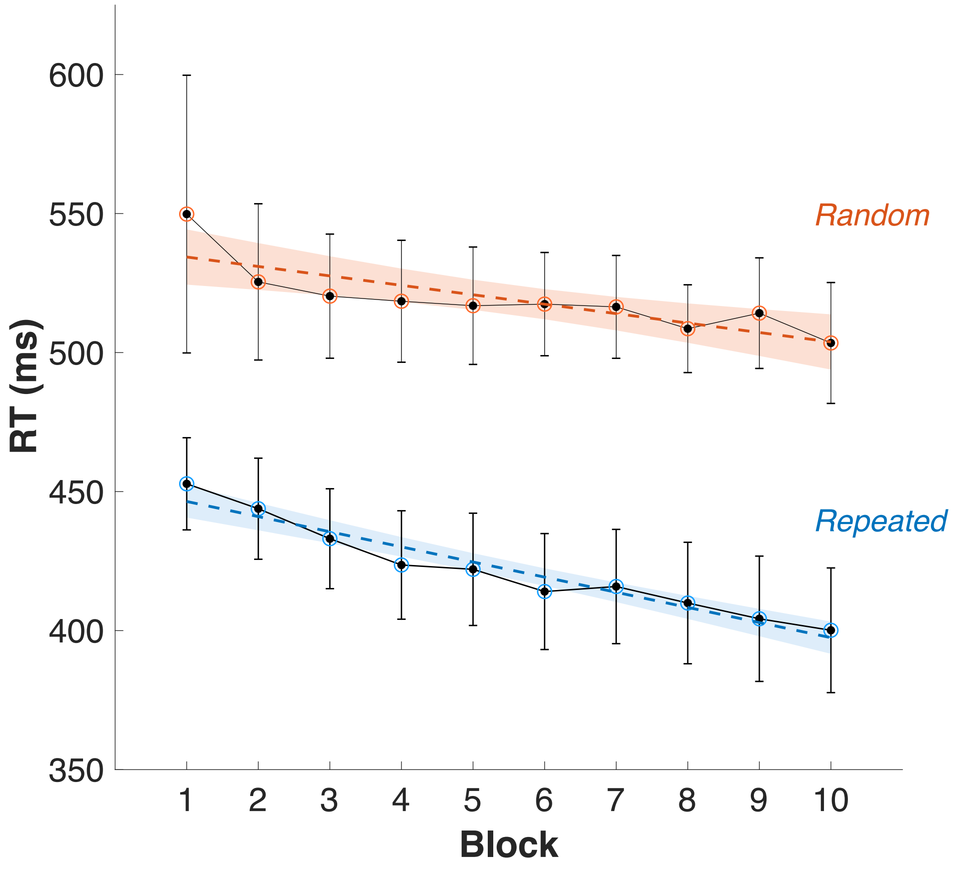


*Note*. A. Mean RTs per block for the baseline acquisition (orange) and mean RTs per block for the experimental task (blue). Results of a multiple linear regression indicated a collective effect of Block, Condition (baseline or experimental) and interaction between predictors ($F$(3,16)=468.4, $p$<.001, R2=.99). The individual predictors were examined further and showed that Block ($t$=-5.14, $p$<.001) and Condition ($t$=-14.82, $p$<.001) are significant predictors in the model, and so is the interaction between them ($t$=-2.19, $p$=.043). This indicates that mean RTs decrease faster in the experimental condition that in the baseline condition, as a result of learning of the repeated sequence and not simply general task learning. Error bars represent 95% confidence intervals.
